# Cortical actin properties controlled by *Drosophila* Fos aid macrophage infiltration against surrounding tissue resistance

**DOI:** 10.1101/2020.09.18.301481

**Authors:** Vera Belyaeva, Stephanie Wachner, Attila Gyoergy, Shamsi Emtenani, Igor Gridchyn, Maria Akhmanova, Markus Linder, Marko Roblek, Maria Sibilia, Daria Siekhaus

## Abstract

The infiltration of immune cells into tissues underlies the establishment of tissue resident macrophages, and responses to infections and tumors. Yet the mechanisms immune cells utilize to negotiate tissue barriers in living organisms are not well understood, and a role for cortical actin has not been examined. Here we find that the tissue invasion of *Drosophila* macrophages, also known as plasmatocytes or hemocytes, utilizes enhanced cortical F-actin levels stimulated by the *Drosophila* member of the fos proto oncogene transcription factor family (Dfos, Kayak). RNA sequencing analysis and live imaging show that Dfos enhances F-actin levels around the entire macrophage surface by increasing mRNA levels of the membrane spanning molecular scaffold tetraspanin TM4SF, and the actin cross-linking filamin Cheerio which are themselves required for invasion. Both the filamin and the tetraspanin enhance the cortical activity of Rho1 and the formin Diaphanous and thus the assembly of cortical actin, which is a critical function since expressing a dominant active form of Diaphanous can rescue the *Dfos* macrophage invasion defect. *In vivo* imaging shows that Dfos enhances the efficiency of the initial phases of macrophage tissue entry. Genetic evidence argues that this Dfos-induced program in macrophages counteracts the constraint produced by the tension of surrounding tissues and buffers the properties of the macrophage nucleus from affecting tissue entry. We thus identify strengthening the cortical actin cytoskeleton through Dfos as a key process allowing efficient forward movement of an immune cell into surrounding tissues.

## Introduction

The classical model of cell migration on a surface postulated in the 1980’s by Abercrombie has been extended (Danuser et al., 2013) by studies showing that migrating cells utilize diverse strategies depending on the architecture and physical properties of their three dimensional (3D) surroundings (Paluch et al., 2016). Much of this work has been conducted *in vitro*, where variations in the environment can be strictly controlled. However most 3D migration occurs within the body, and much less research has elucidated the mechanisms used to efficiently move in these diverse environments, particularly into and through tissues. Such migration is crucial for the influence of the immune system on health and disease. Vertebrate macrophages migrate into tissues during development where they take up residence, regulating organ formation and homeostasis and organizing tissue repair upon injury (Ginhoux and Guilliams, 2016; Theret et al 2019). A variety of types of immune cells infiltrate into tumors, and can both promote or impede cancer progression (Greten and Grivennikov 2019; Sharma and Allison, 2015). Responses to infection require immune cells to traverse through the vascular wall, into the lymph node, and through tissues (Luster et al., 2005). Yet the mechanisms utilized by immune cells to allow migration into such challenging cellular environments *in vivo* are not well understood.

Migration in 2D and 3D environments requires actin polymerization to power forward progress. The assembly of actin at the leading edge, when coupled to Integrin adhesion to anchor points in the surrounding ECM, can allow the front of the cell to progress (Mitchison and Cramer, 1996). This anchoring also allows the contraction of cortical actin at the rear plasma membrane to bring the body of the cell forwards. But a role for crosslinked actin at the cell surface in assisting forward progress by helping to counteract the resistance of surrounding tissues and in buffering the nucleus has not been previously identified.

Our lab examines *Drosophila* macrophage migration into the embryonic germband (gb) to investigate mechanisms of immune cell tissue invasion. Macrophages, also called plasmatocytes or hemocytes, are the primary phagocytic cell in *Drosophila* and share striking similarities with vertebrate macrophages (Brückner et al., 2004; Evans & Wood, 2011; Lemaitre & Hoffmann, 2007; Ratheesh et al., 2015; Weavers et al., 2016). They are specified in the head mesoderm at embryonic stages 4-6 and by stage 10 start spreading along predetermined routes guided by platelet-derived growth factor- and vascular endothelial growth factor-related factors (Pvf) 2 and 3 (Cho et al., 2002; Brückner et al., 2004; Wood et al., 2006) to populate the whole embryo. One of these paths, the movement into the gb, requires macrophages to invade confined between the ectoderm and mesoderm (Ratheesh et al., 2018; Siekhaus et al., 2010). The level of tension and thus apparent stiffness of the flanking ectoderm is a key parameter defining the efficiency of macrophage passage into and within the gb (Ratheesh et al., 2018). Penetration of macrophages into the gb utilizes Integrin, occurs normally without MMPs (Siekhaus et al., 2010) and is even enhanced by ECM deposition (Valoskova et al., 2019; Sánchez-Sánchez et al., 2017) likely because the basement membrane has not yet formed at this stage (Matsubayashi et al., 2017; Ratheesh et al., 2018). Thus *Drosophila* macrophage gb invasion represents an ideal system to explore the mechanisms by which immune cells and surrounding tissues interact with one another to aid the invasion process.

Here we sought to identify a transcription factor that could control immune cell tissue invasion and elucidate its downstream mechanisms. We identify a role for the *Drosophila* ortholog of the proto-oncogene Fos in initial entry and migration within the tissue. We find Dfos increases cortical macrophage F-actin levels through the formin Cheerio and a novel target, the tetraspanin TM4SF, aiding macrophages to move forward against the resistance of the surrounding tissues while buffering the nucleus.

## Results

### The transcription factor Dfos is required for macrophage germband invasion

To identify regulators of programs for invasion we searched the literature for transcription factors expressed in macrophages prior to or during their invasion of germband tissues (gb) (Fig 1A-B’). Of the 12 such factors (S1 Table, based on Hammonds et al., 2013) we focused on Dfos, a member of the Fos proto-oncogene family, assigned by the Roundup algorithm as being closest to vertebrate c-fos (Deluca et al., 2012; Thurmond et al., 2019) (Fig 1C). Dfos contains the basic leucine zipper domain (bZIP) shown to mediate DNA binding and hetero and homo dimerization (Glover and Harrison, 1995; Szalóki et al., 2015) with the third leucine replaced by a methionine, a position also altered in the *C. elegans* ortholog FOS-1A (Sherwood et al., 2005). Embryo *in situ* hybridizations reveal enriched expression of the gene in macrophages at early stage 11 (Fig 1D, arrow) which is attenuated by stage 13 matching what was seen in the BDGP *in situ* database https://insitu.fruitfly.org/cgi-bin/ex/report.pl?ftype=1&ftext=FBgn0001297. Antibody staining against Dfos protein appears in the nucleus in macrophages that are migrating towards the gb at stage 10-12 (Fig 1E-F’ yellow arrowheads, G-G”’ white arrows) and is still observed in stage 13 (S1A Fig). The *Dfos^1^* null mutant that removes exon 1 including the translational start site (Riesgo-Escover and Hafen, 1997, Zeitlinger, et al., 1997) eliminates the signal in macrophages, indicating antibody specificity (Fig 1H). To determine if Dfos affects invasion, we examined the 70% of embryos that did not display developmental defects at these early stages from *Dfos^1^* and the hypomorphic *Dfos^2^* (Zeitlinger et al., 1997); we quantified macrophage numbers in the gb during a defined developmental period in early stage 12 (Fig 1M). Both Dfos mutants displayed significantly reduced numbers of macrophages in the gb compared to the control (Fig 1I-K, N) with normal numbers in the pre-gb zone for *Dfos^2^* (S1B Fig) (S1 Data). Macrophage-specific expression of *Dfos* rescues the *Dfos^2^* mutant (Fig 1L,N). Blocking Dfos function in macrophages with a dominant negative (DN) Dfos (Fig 1O-Q) that lacks the activation domain but retains the capacity to dimerize and bind DNA (Eresh et al., 1997) or two different RNAis against *Dfos* (Fig 1R) recapitulates the decrease in gb macrophages seen in the null while not affecting macrophage numbers in the whole embryo (S1C Fig), and along the ventral nerve cord (vnc) (S1D-E Fig). However, macrophages expressing DfosDN or the *Dfos* RNAis accumulate in the pre-germband area (S1F-G Fig), as if they are accumulating there when unable to progress further. These results argue that Dfos is required in macrophages for their migration into the gb. The tool we chose to examine this capability was DfosDN for the following reasons. Dfos and DfosDN do not appear to inhibit other bZIP proteins at higher levels of expression: overexpressing DfosDN in the midgut does not inhibit another bZIP protein that acts there (Eresh et al., 1997) and overexpressing Dfos in macrophages does not change gb numbers (S1H Fig). DfosDN should exert a quicker effect than the RNAis. And finally, the *Dfos RNAi*s no longer exert an effect when a second UAS construct is simultaneously expressed (S1I Fig). Thus our further experiments examining Dfos’ role in enhancing macrophage germband invasion utilized mostly the DN form.

**Fig 1.**
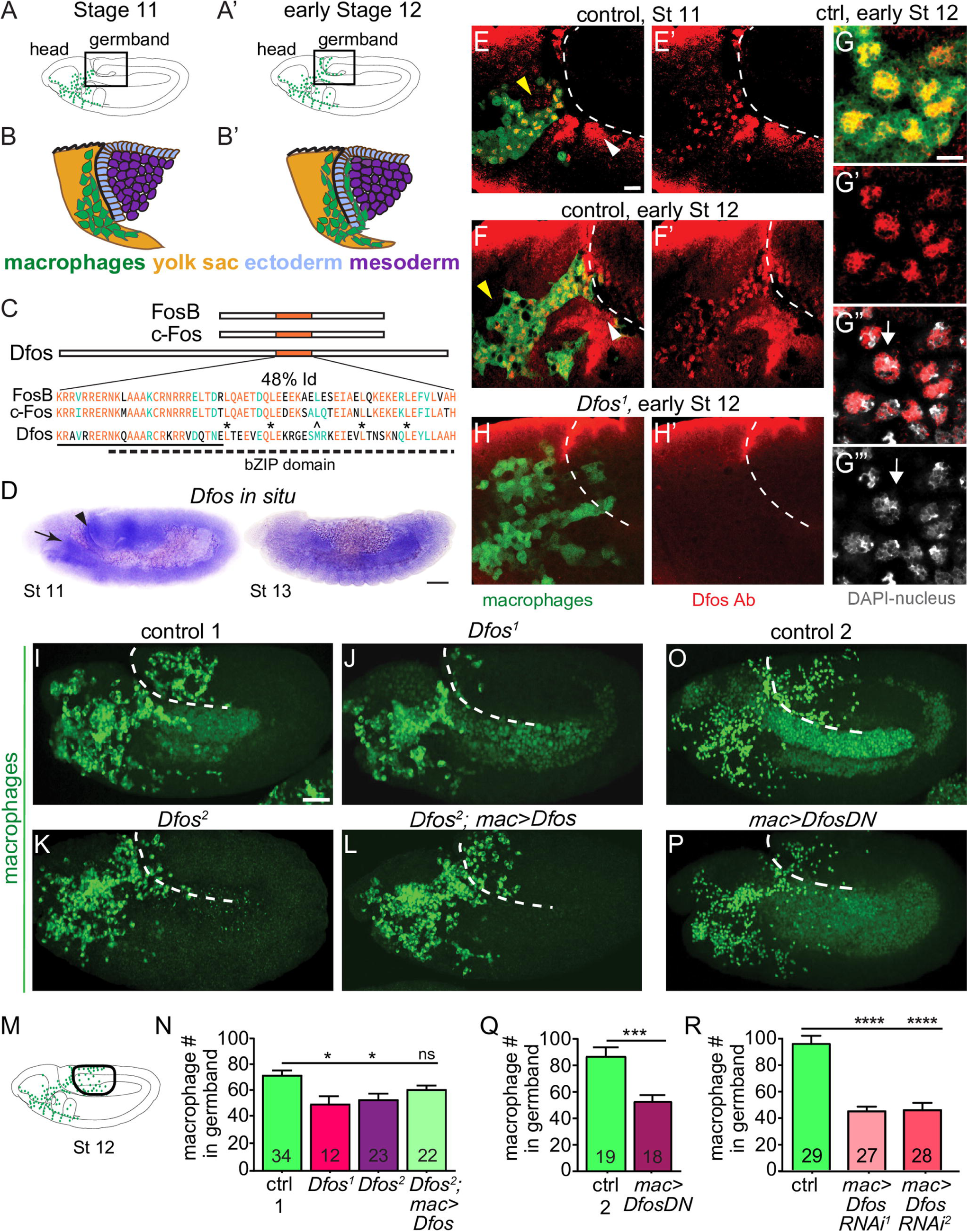
The bZIP transcription factor Dfos acts in macrophages to facilitate their migration into the germband. Schematics of lateral **(A)** stage (St) 11 and **(A’)** early St 12 embryos. The boxed region magnified below indicates where macrophages (green) invade the germband (gb) after moving there from the head **(B-B’)**. Macrophages sit on the yolk sac (yellow) next to the amnioserosa (black line) and then invade between the ectoderm (blue) and mesoderm (purple). (**C**) Dfos protein aligned with its human orthologs c-Fos and FosB; orange outlines the bZIP region that has 48% identity to both proteins: identical amino acids shown in orange, conserved ones in green. Stars indicate Leucines in the zipper; ^ the third leucine which in Dfos is a methionine, a tolerated substitution (Garcia-Echeverria, 1997). The lower solid line indicates the basic domain and the dotted line the leucine zipper (ZIP). (**D**) *In situ* hybridization of St 11 and 13 embryos with a riboprobe for Dfos-RB (Fbcl0282531) which also detects all Dfos isoforms. *Dfos* RNA expression is enriched in macrophages (arrow) and the amnioserosa (arrowhead) before gb invasion, but is gone thereafter. **(E-H’)** Confocal images of the boxed region in A from fixed embryos expressing *GFP* in macrophages (green) stained with a Dfos Ab (red). **(E-F’, H-H’)** A white dashed line indicates the gb edge. **(E-F)** The Dfos Ab (yellow arrowheads) stains (**E**) macrophages moving towards the gb at St 11, and **(F)** early St 12, as well as the amnioserosa (white arrowheads). **(G)** Higher magnification shows Dfos colocalizing with the nuclear marker DAPI (white). **(H)** No staining is detected in macrophages or the amnioserosa in the null *Dfos^1^* mutant. **(I-L)** Lateral views of mid St 12 embryos from **(I)** the control, **(J)** the null allele *Dfos^1^*, **(K)** the hypomorphic allele *Dfos^2^*, and **(L)** *Dfos^2^* with *Dfos* re-expressed in macrophages. **(M)** Schematic of St 12 embryo, gb region indicated by a black oval outline. **(N)** Quantitation reveals that both *Dfos* alleles display fewer macrophages in the gb. Re-expression of *Dfos* in macrophages in the *Dfos^2^* hypomorph significantly rescues the defect. Control vs. *Dfos^1^* p=0.02 (30% reduction), Control vs. *Dfos^2^* p=0.017 (25% reduction), Control vs. *Dfos^2^; mac>Dfos* p=0.334. **(O-P)** Lateral views of mid St 12 embryos from **(O)** the control, or **(P)** a line expressing a dominant negative (DN) form of Dfos in macrophages. **(Q)** Quantification of macrophage numbers in the gb (see schematic) in the two genotypes visualized in **O**-**P**. p=0.0002 (***) (40% reduction). Standard Deviation (SD): 25, 25. **(R)** Quantification of macrophage numbers in the gb of the control and two different lines expressing RNAi constructs against Dfos in macrophages. Quantification of macrophage numbers in the gb for lines expressing one of two different *UAS-Dfos RNAi* constructs in macrophages. Control vs. *mac>Dfos RNAi^1^* (TRiP HMS00254) or vs. *mac>Dfos RNAi^2^* (TRiP JF02804), p<0.0001 (54 or 52% reduction). SD: 32, 19, 29. The data in **Q** and **R** argue that Dfos is required within macrophages to promote gb tissue invasion. Embryos are positioned with anterior to left and dorsal up in all images and histograms show mean + standard error of the mean (SEM) throughout. Macrophages are labeled using *srp-Gal4* (“mac>”) driving *UAS-GFP* in **E-H**, *UAS-GFP::nls* in **I-L** and *srpHemo-H2A::3xmCherry* in **O-R**. ***p<0.005, **p<0.01, *p<0.05. One-way ANOVA with Tukey post hoc was used for **N** and **R**, and unpaired t-test for **Q**. The embryo number analysed is indicated within the relevant column in the graphs. Scale bar: 50 µm in **D**, 5 µm in **E-H** and 10 µm in **I-L, O-P**.

### Dfos promotes macrophage motility and persistence during tissue entry

To examine the dynamic effects of Dfos on tissue invasion, we performed live imaging and tracking of macrophages. We visualized macrophages with *srpHemo-H2A::3xmCherry* (Gyoergy et al., 2018) in either a wild type or *mac>DfosDN* background, capturing the initial stage of invasion (S1 Movie). The speed of macrophages moving in the area neighboring the germband prior to invasion was not significantly changed (pre-gb, Fig 2B,C). However, the first *mac>DfosDN* macrophage to enter is delayed by 20 min in crossing into the gb (Fig 2D). *mac>DfosDN* macrophages also displayed reduced speed and directional persistence during entering as well as while moving along the first 20μm of the ectoderm-mesoderm interface (gb entry, Fig 2E, S2A Fig). Macrophages in the *Dfos^2^* mutant largely mirrored this phenotype, but displayed slower movement in the pre-gb zone neighboring the amnioserosa in which Dfos is also expressed (Fig 1D-F), likely causing a non-autonomous effect (S2B-C Fig, S2 Movie) (Fig 1D, black arrowhead, E-F, white arrowheads). Macrophages expressing DfosDN moved with unaltered average speed as they spread out along the non-invasive route of the vnc (Fig 2F, Fig 2G, S3 Movie), albeit with reduced directional persistence (S2A Fig). We thus conclude from live imaging that Dfos in macrophages aids their initial invasive migration into the gb, increases their speed within the gb and does not underlie their progress along the vnc.

**Fig 2.**
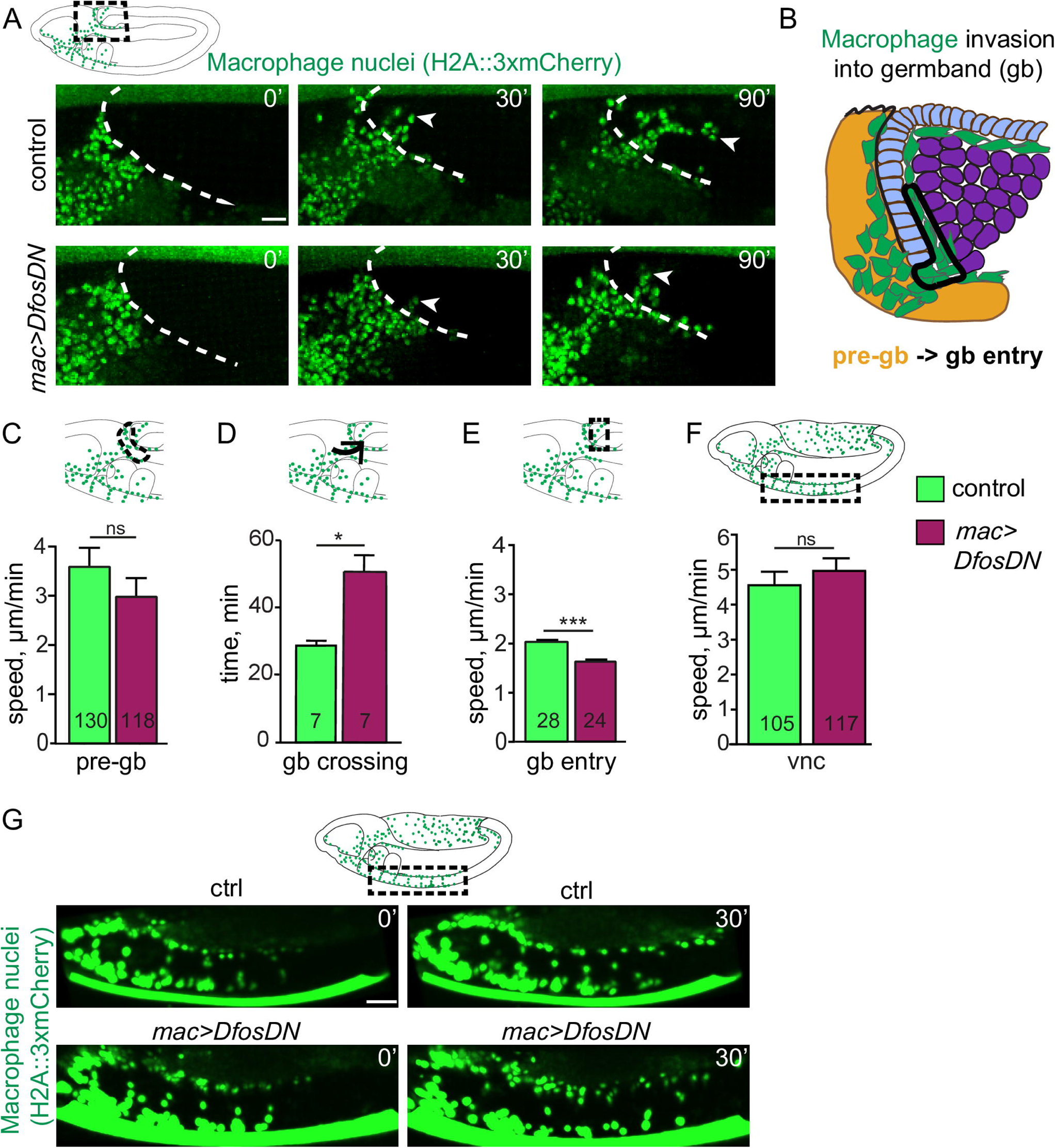
Dfos facilitates the initial invasion of macrophages into the gb tissue. (**A**) Movie stills of control embryos and those expressing DfosDN in macrophages (green, labelled using *srpHemo-H2A::3xmCherry*). Area imaged corresponds to the black dashed square in the schematic above. The germband (gb) border is outlined with a white dashed line. The first entering macrophage is indicated with a white arrowhead, and time in minutes in the upper right corner. (**B**) Detailed schematic showing the different zones for which the parameters of macrophage gb invasion were quantified. The pre-gb area is shown in yellow, the gb entry zone is outlined in a solid line. (**C**) Macrophage speed in the pre-gb area was not significantly changed in macrophages expressing DfosDN (3.00 µm/min) compared to the control (3.61 µm/min), p= 0.58. (**D**) Quantification shows a 68% increase in the total gb crossing time of DfosDN expressing macrophages compared to the control. Total gb crossing time runs from when macrophages have migrated onto the outer edge of the gb ectoderm, aligning in a half arch, until the first macrophage has translocated its nucleus into the gb ecto-meso interface. p=0.008. SD: 4, 14. (**E**) DfosDN expressing macrophages displayed a significantly reduced speed (1.53 µm/min) at the gb entry zone compared to the control (1.98 µm/min), p= 1.11e^-06^. SD: 2, 2. (**F**) Macrophages expressing DfosDN in a Stage 13 embryo move with unaltered speed along the vnc in the region outlined by the dashed black box in the schematic above (4.93 µm/min), compared to the control (4.55 µm/min), p= 0.64. Corresponding stills shown in **(G**) Macrophages are labeled by *srpHemo-Gal4* driving *UAS-GFP::nls*. ***p<0.005, **p<0.01, *p<0.05. Unpaired t-test used for **C-F**, a Kolmogorov-Smirnov test for **D.** For each genotype, the number of tracks analysed in **C** and **F,** and the number of macrophages in **D-E** are indicated within the graph columns. Tracks were obtained from movies of 7 control and 7 *mac>DfosDN* expressing embryos in panel **D**, 3 each in **C, F**, and 4 each in **E**. Scale bar: 10 µm.

### Dfos modulates Filamin and Tetraspanin to aid gb tissue invasion

To identify Dfos targets that promote macrophage invasion, we FACS isolated macrophages from wild type and *mac>DfosDN* embryos during the time when invasion has just begun, and conducted RNA-sequencing of the corresponding transcriptomes (Fig 3A, S1 Data). We first assessed reads that map to Dfos, which can correspond to both endogenous and DfosDN mRNA; we found a 1.6 fold increase in the presence of the one copy of DfosDN in this line, arguing that this transgene is expressed at levels similar to each endogenous copy of Dfos and is unlikely to produce extraneous effects (S2 Data). We then examined genes that displayed a log2 fold change of at least 1.5 with an adjusted P value less than 0.05 in the presence of DfosDN. Ten genes were down-regulated (Fig 3B, S3A-B Fig) and 9 up-regulated by DfosDN (S2 Table). Upregulated genes in DfosDN encoded mostly stress response proteins, consistent with the role previously demonstrated for fos in *C. elegans* in suppressing stress responses (Hattori et al., 2013). We concentrated on the downregulated class. Of these, we focused on the actin crosslinking filamin Cheerio (Cher) and the tetraspanin TM4SF from a group that can form membrane microdomains that affect signalling and migration (Razinia et al., 2012; Yeung et al., 2018). No known role for TM4SF had been previously identified in *Drosophila*. To determine if these Dfos targets were themselves required for invasion, we knocked down Cher and TM4SF through RNAi individually or simultaneously and observed significantly reduced macrophage numbers in the gb, particularly upon the knockdown of both targets simultaneously (Fig 3C-G) while not affecting macrophage numbers in the pre-gb zone (S3D Fig) or on the vnc (S3E Fig). Over-expression of Cher or TM4SF along with *DfosDN* in macrophages increased the mean macrophage numbers in the gb, and over-expression of TM4SF rescued the *DfosDN* macrophage invasion defect (Fig 3H-L). Expression of a GFP control did not restore macrophage invasion indicating that the rescue we observed through Cher or TM4SF expression was not due to promoter competition leading to reductions in DfosDN expression. We conclude that Dfos aids macrophage gb invasion by increasing the mRNA levels of the filamin actin crosslinker Cher and the tetraspanin TM4SF.

**Fig 3.**
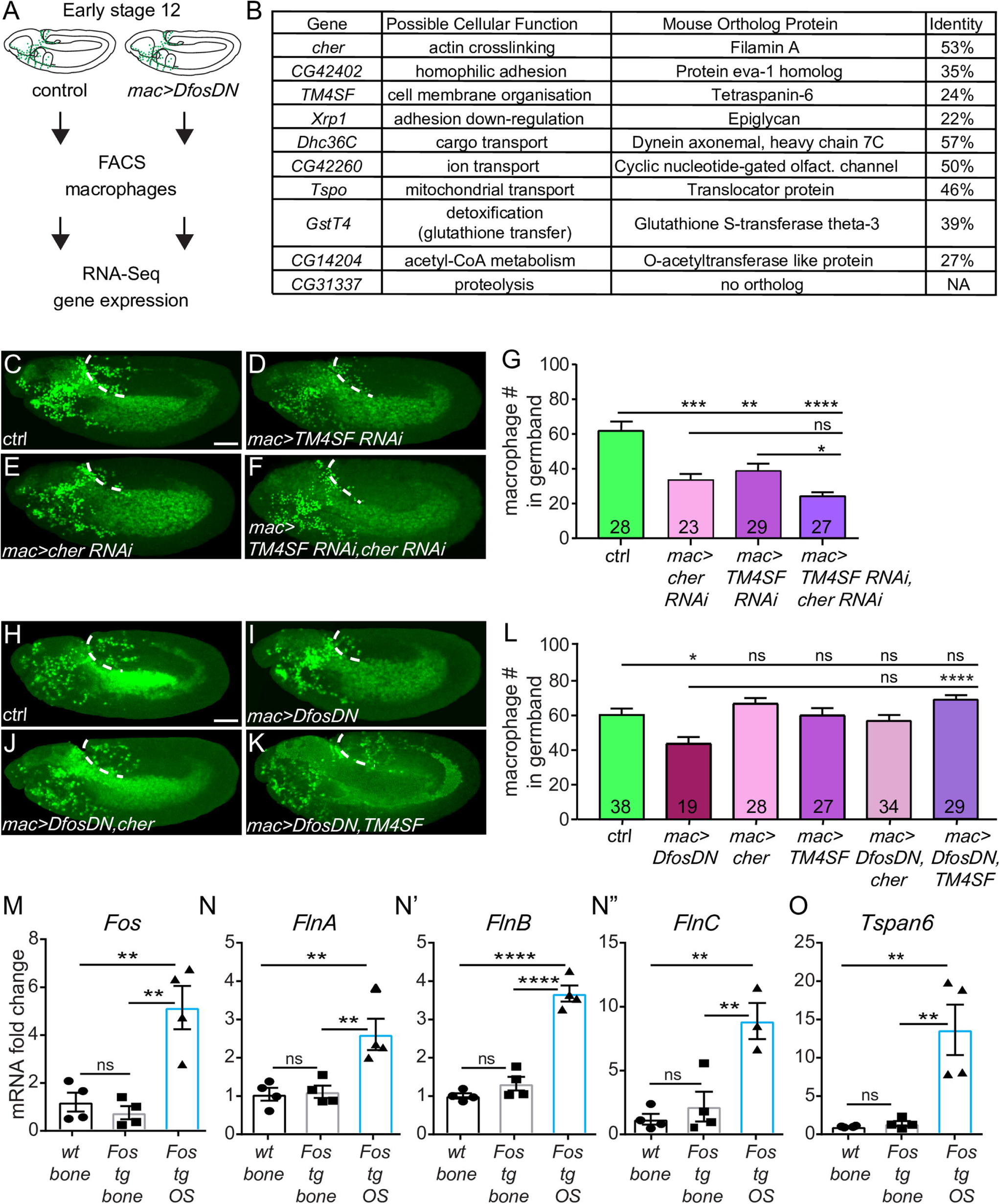
Dfos regulates macrophage germband invasion through cytoskeletal regulators: the Filamin Cheerio and the tetraspanin TM4SF. **(A)** Schematic representing the pipeline for analyzing mRNA levels in FACS sorted macrophages. **(B)** Table of genes down-regulated in macrophages expressing DfosDN. Genes are ordered according to the normalized p-value from the RNA-Sequencing. The closest mouse protein orthologs were found using UniProt BLAST; the hit with the top score is shown in the table. **(C-F)** Lateral views of representative St 12 embryos in which the two targets with links to actin organization, (**D**) the Tetraspanin TM4SF and (**E**) the Filamin Cheerio, have been knocked down individually or (**F**) together, along with the control (**C**). Scale bar: 50 μm. **(G)** Quantification shows that the number of macrophages in the germband is reduced in embryos expressing RNAi against either *cher (KK 107451*) or *TM4SF* (*KK 102206*) in macrophages, and even more strongly affected in the double RNAi of both. Control vs. *cher RNAi* p=0.0005 (46% reduction). Control vs. *TM4SF RNAi* p=0.009 (37% reduction), Control vs. *cher/TM4SF RNAi* p>0.0001 (61% reduction). *cher RNAi* vs. *TM4SF RNAi* p=0.15. SD: 29, 23, 17, 12. **(H-K)** Lateral views of a representative St 12 embryo from (**H**) the control, as well as embryos expressing DfosDN in macrophages along with either (**I**) GFP, (**J**) Cher, or (**K**) TM4SF. (**L**) Quantification shows that over-expression of TM4SF in DfosDN expressing macrophages restores their normal numbers in the gb. Over-expression of Cher in this background shows a strong trend towards rescue, but did not reach statistical significance. Control vs. *DfosDN* p=0.015 (28% reduction); Control vs. *cher* p=0.74; Control vs. *TM4SF* p>0.99; *DfosDN* vs. *DfosDN cher* p=0.14; DfosDN vs. *DfosDN, TM4SF* p<0.0001; Control vs. *cher* p=0.97; Control vs. *TM4SF* p=0.35. SD: 22, 16, 16, 21, 22, 13. **(M-O)** q-PCR analysis of mRNA extracted from the bones of mice that are wild type, transgenic (tg) for *Fos* controlled by a Major Histocompatibility promoter and viral 3’UTR elements, and those in which such c-Fos transgenesis has led to an osteosarcoma (OS). Analysis of mRNA expression shows that higher levels of (M) *Fos* correlate with higher levels of **(N-N”)** FlnA-C, and **(O)** Tspan6 in osteosarcomas. p values = 0.86, 0.001, 0.003, SD: 0.7, 0.6, 0.3 in **M**, 0.98, 0.009, 0.007 and 0.4, 0.2, 1.5 in **N**, 0.39, < 0.0001, <0.0001 and 0.2, 0,3, 1.1 in **N’**, 0.76, 0.005, 0.002 and 0.8, 2.3, 2.4 in **N”**, 0.99, 0.004, 0.003 and 0.1, 0.2, 0.2 in **O**. Scale bar: 50 μm. Macrophages are labeled using either (C-F) *srp::H2A::3xmCherry* or (H-K) *srpHemo-Gal4* (“*mac>*”) driving *UAS-mCherry::nls*. ***p<0.005, **p<0.01, *p<0.05. Unpaired t-test or one-way ANOVA with Tukey post hoc were used for statistics. Each column contains the number of analyzed embryos.

### In murine osteosarcoma c-fos mRNA level increases correlate with those of Filamins and Tetraspanin-6

To determine if these Dfos targets in *Drosophila* could also be Fos targets in vertebrate cells, we utilized a well-established murine transgenic model that over expresses c-fos. In these mice transgenic c-fos expression from viral 3’ UTR elements in osteoblasts (the bone forming cells) leads to osteosarcoma development accompanied by a 5 fold increase in c-fos mRNA expression (Fig 3M) (Linder et al., 2018). We examined by qPCR the mRNA levels of our identified Dfos targets’ orthologs, comparing their levels in osteosarcomas (Fos tg OS) to neighboring, osteoblast-containing healthy bones from Fos tg mice (Fos tg bone) and control bones from wild-type mice (wt bone). We saw 2.5 to 8 fold higher mRNA levels of the three murine Filamin orthologs (Fig 3N-N”) and a 15 fold increase in Tetraspanin-6 (Fig 3O) in osteosarcoma cells. mRNA levels of several of the orthologs of other Dfos targets we had identified showed less strong inductions or even decreases; the Glutathione S transferase Gstt3 and the Slit receptor Eva1c increased 4 and 2.8 fold respectively, while the mitochondrial translocator Tspo was 25% lower (S3F-I Fig). These results suggest that Dfos’s ability to increase mRNA levels of two key functional targets for migration, a Filamin and a Tetraspanin, is maintained by at least one vertebrate fos family member.

### Dfos increases assembly of cortical actin through Cheerio and TM4SF to aid macrophage invasion

We wished to determine what cellular properties Dfos could affect through such targets to facilitate *Drosophila* macrophage invasion. Given Cheerio’s known role as an actin crosslinker, we examined actin in invading *mac>DfosDN* macrophages within live embryos. To visualize actin in macrophages, we utilized a *srpHemo-moe::3xmCherry* reporter which marks cortical F-actin (Edwards et al., 1997; Franck et al., 1993) and observed a reduction of 53% (Fig 4A-D) in its signal in invading m*ac>DfosDN* macrophages. We saw no change by Western analysis in the levels of the Moe::3xmCherry protein itself upon DfosDN expression (Fig S4A-A’). We hypothesized that the changes in cortical actin we observed in the *mac>DfosDN* all could be due to the lower levels of Cheerio and/or TM4SF mRNA. Indeed, we observed reductions in Moe::3xmCherry all around the edge of invading macrophages in live embryos expressing RNAi against Cher or TM4SF in macrophages (Fig 4E-H). To test if a decrease in actin assembly could underlie the reduced tissue invasion of *mac>DfosDN* macrophages, we forced cortical actin polymerization by expressing a constitutively active version of the formin Diaphanous (DiaCA) which localizes to the cortex (Gonzalez-Gaitan and Peifer, 2009). Indeed, expressing DiaCA in macrophages completely rescued the *Dfos^1^*, *Dfos^2^* (Fig S4B), and *mac>DfosDN* invasion defect (Fig 4I-J). Given that Dia, like Dfos, does not affect general macrophage migratory capacities along the ventral nerve cord (Davis et al., 2015), we examined if Dia might normally play a role in invasion. We utilized two RNAis against Dia and observed decreased macrophage numbers in the gb in each (Fig 4K-L) with no effect on numbers in the pre-gb (S4C Fig) or on the vnc (S4D Fig). These results argue that Dfos aids invasion by increasing levels of TM4SF and Cheerio to enhance assembly of actin around the surface of the macrophage, potentially by increasing the activity of Dia.

**Fig 4.**
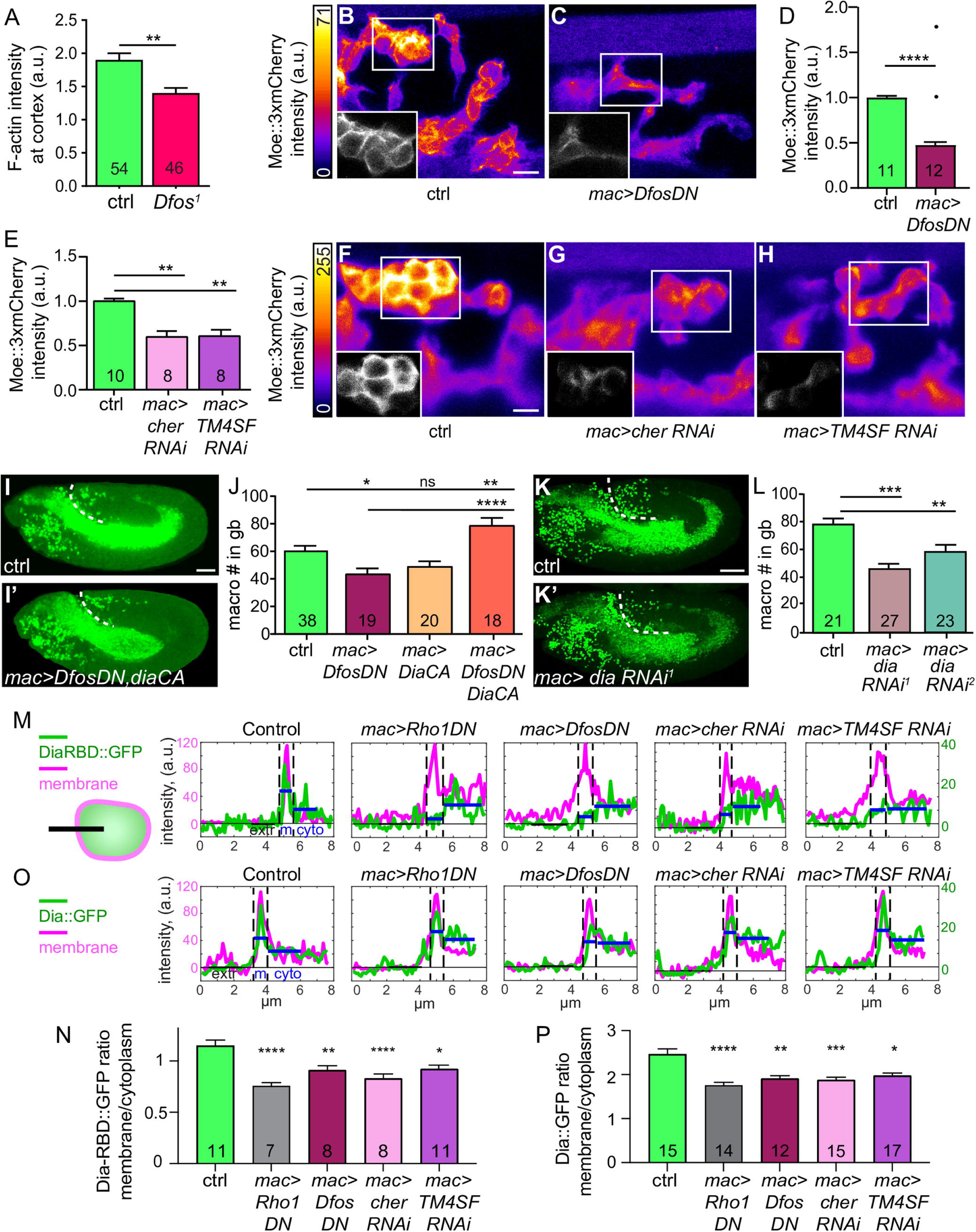
Dfos regulates the actin cytoskeleton through Cher, TM4SF, and the formin Diaphanous. **(A)** Quantification of phalloidin intensity to detect F actin at the macrophage-macrophage contacts in Stage 11/12 *Dfos^1^* embryos. F-actin is strongly reduced at these homotypic contacts. **(B-C)** Representative confocal images of live embryos expressing in invading macrophages the F-actin binding and homodimerizing portion of Moesin (*srpHemo-moe::3xmCherry)* to label F-actin, presented as a maximum z-projection. Relative Moe-3xmCherry intensity is indicated with a pseudo-color heat map as indicated on the left, with yellow as the highest levels and dark blue as the lowest as indicated in the calibration bar to the left. Insets in the bottom left corner of each panel show a grey-scale single z-plane corresponding to the white box in the main image. Embryo genotype indicated below. Strong reductions in cortical actin are observed in macrophages expressing DfosDN compared to the control. **(D-E)** Quantification of the macrophage Moe:3xmCherry intensity as a measure of cortical F-actin, normalized to the average fluorescence intensity of the control per batch. **(D)** Quantification shows that macrophages expressing DfosDN display a 53% reduction in Moe::3xmCherry intensity compared to the control when the two outliers shown as single dots are excluded, 37% if they are included. Outliers identified by 10% ROUT. n of ROIs analysed = 650 for control, 687 for *DfosDN*. p=0.0007 for analysis including outliers (Kolmogorov-Smirnov) and p<0.0001 for analysis excluding outliers (Welch’s t-test). SD: 0.2, 0.4. **(E)** Quantification reveals that macrophage expression of an RNAi against either *cher* or *TM4SF*, the two genes whose expression is reduced in *DfosDN*, also results in a decrease of Moe::3xmCherry intensity (by 40% each). n of ROIs analysed = 549 for control, 423 for *cher RNAi,* 306 for *TM4SF RNAi*. Control vs. *cher RNAi* p=0.006. Control vs. *TM4SF* p=0.003. SD: 0.2, 0.3, 0.2. **(F-H)** Images and representation as in B-C. Strong reductions in cortical actin are observed in macrophages expressing *cher RNAi* or *TM4SF RNAi* compared to the control. **(I,I’)** Representative confocal images of St 12 embryos from the control and a line in which macrophages express DfosDN and a constitutively active (CA) form of the formin Dia to restore cortical actin polymerization. **(J)** Quantification shows that while macrophage expression of DiaCA does not significantly affect the number of macrophages in the gb, expressing it in a DfosDN background rescues macrophage gb invasion. Control vs. *DfosDN* p=0.017 (28% reduction), Control vs. *diaCA* p=0.18, Control vs. *DfosDN*, *diaCA* p=0.010, *DfosDN* vs. *DfosDN, diaCA* p<0.0001. SD: 22, 16, 16, 24. (**K,K’**) Representative confocal images of St 12 embryos from the control and from a line expressing an RNAi against *dia* in macrophages. **(L)** Quantification of two RNAi lines against *dia* expressed in macrophages shows a 37% and 21% reduction in macrophage numbers in the gb compared to control. Control vs. *dia RNAi^1^* (TRiP HMS05027) p<0. 0001; control vs. *dia RNAi^2^* (TRiP HMS00308) p=0.0008. SD: 13, 20, 22. (**M, O**) Examples of line profiles used for the determination of the membrane-to-cytoplasmic ratio of DiaRBD in panel N and Dia in panel P. Line intensity profiles from fixed Stage 11 embryos of (M) DiaRBD::GFP or (O) Dia::GFP (green) and membrane myr::Tomato (magenta) across the outward facing edge of groups of macrophages sitting within ~ 40 µm of the germband that expressed either lacZ (Control), Rho1DN, DfosDN, *cher RNAi*, or *TM4SF RNAi* as shown in the schematic in M. Line length ~ 8 µm. Blue lines indicate mean GFP intensity on the membrane and in cytoplasm. (**N, P**) Quantification of membrane-to-cytoplasmic intensity ratio of (N) DiaRBD::GFP or (P) Dia::GFP expressed in macrophages under UAS control along with either lacZ (control, n=158 line scans from 11 embryos or n=233 from 15), Rho1DN (n=123 from 7 or n=212 from 14), DfosDN (n=135 from 8 or n=237 from 12), *cher RNAi* (n=128 from 8 or n=252 from 13), *TM4SF RNAi* (n=205 from 11 or n=279 from 17). Control vs. Rho1DN ****p<0.0001 (34% or 29% reduction), Control vs. DfosDN p=**0.0037 (21 or 23% reduction), Control vs. *cher RNAi* ***p=0.0007 (28 or 24% reduction), Control vs. *TM4SF RNAi* *p=0.026 or 0.024 (20% reduction). SD: 0.7, 0.5, 0.5, 0.5, 0.4 in N; 1.9, 0.9, 1.0, 0.9, 1.0 in P. Macrophages are labeled using either *srpHemo-Gal4* driving *UAS-mCherry::nls* **(I-I’)**, *srpHemo-H2A::3xmCherry* **(K-K’).** *srpHemo-moe::3xmCherry*, *srpHemo-Gal4* (*mac>)* crossed to **(B)** *UAS-GFP* as a Control, **(C)** *UAS-DfosDN*, **(F)** *w^-^* Control, **(G)** *UAS-cher RNAi* (KK 107451), **(H)** *UAS-TM4SF RNAi* (KK 102206). *srpHemo-GAL4 UAS-Myr::tdTomato UAS-diaRBD::GFP* (M, O) *or UAS-dia::GFP* (N, P) crossed to *UAS-lacZ* as a Ctrl, *UAS-Rho1DN* or the lines indicated above. ***p<0.005, **p<0.01, *p<0.05. Unpaired t-test used for **A.** Welch’s t test of normalized average mean intensity per embryo for **D** with the two indicated outliers excluded, for statistical assessment. One way ANOVA with Tukey post hoc for **E**, **J, L.** Kruskal-Wallis for **N, P**. The number of analyzed **(A)** macrophage-macrophage junctions, or **(D-E, J, L, N, P)** embryos is shown in each column. Scale bar 10 μm in **(B-C, F-H),** 50 μm in (**I, K).**

### Dfos stimulates the cortical activity of Rho1 and Diaphanous through its targets TM4SF and Cheerio

We sought to examine if Dfos or its targets could stimulate Dia activity or that of its known regulator, Rho, at the cortex. Dia’s autoinhibition negatively regulates its cortical localization and activity (Goode, Eck Ann Rev Biochemistry, 2007). For mDia, binding to activated Rho GTPases as well as to other unknown membrane associated proteins can release this autoinhibition (Seth, Otsomo, Rosen JCB 2006). We expressed a reporter for active Rho1, the Rho1 binding domain of Dia (DiaRBD::GFP) (Abreu-Blanco et al., 2014), in macrophages along with myristoylated Tomato to delineate the plasma membrane and quantified intensity profiles of linescans across the membrane in various genetic backgrounds (Fig 4M). To validate the assay we expressed Rho1DN and found, as expected, a significant reduction, by 34%, in cortical DiaRBD compared to the control (mem/cyto=1.15 in control, 0.76 for Rho1DN) (Fig 4N). Interestingly expressing either DfosDN, or RNAis against Cher or TM4SF also resulted in a reduction of cortical DiaRBD, by 21, 28, and 20%, respectively (mem/cyto=0.91, 0.83, 0.92 respectively). We then tested if this increase in Rho1 activity prompted by Dfos and its targets also resulted in a change in the activity and thus the localization of Dia itself, utilizing Dia::GFP (Homem & Peifer, 2008) expressed in macrophages along with Myr::Tomato to mark the membrane and quantification as above (Fig 4O). As predicted by the prior results with mDia, upon the expression of Rho1DN we observed a significant reduction, by 29%, in cortical Dia compared to the control (mem/cyto=2.46 in control, 1.76 for Rho1DN) (Fig 4P). Consistent with our findings for their effect on Rho1 activity, we observed that expressing either DfosDN, or RNAis against Cher or TM4SF resulted in a reduction of cortical Dia, by 23, 23, and 20%, respectively (mem/cyto=1.9, 1.88, 1.97 respectively). Our data argue that the Dfos targets TM4SF and Cheerio increase Rho1’s activity and Dia’s localization at the cortex, stimulating cortical actin assembly.

We examined what consequence these lower cortical F-actin levels had on the cellular behavior of macrophages during entry. Quantitation showed that the actin protrusion that macrophages initially insert between the ectoderm and mesoderm during invasion was actually longer in the *mac>DfosDN >LifeAct::GFP* macrophages than in the control (Fig 5A, S5A Fig, S4 Movie). We then performed live imaging of macrophages labeled with CLIP::GFP to visualize microtubules and thus cell outlines in both genotypes; we determined the aspect ratio (maximal length over width) that the first entering cell displays as it enters into the gb. The first DfosDN-expressing macrophage was extended even before it had fully moved its rear into the gb (S5B Fig). We carried out measurements, taking only cells that had entered the gb to be able to clearly distinguish the rear of the first macrophage from the tips of following cells (Fig 5B). We also avoided including in this measurement the forward protrusion and determined that the first macrophage inside the gb displays an average increase of 23% in the maximal length (L) of the cell body and a 12% reduction in the maximal width (W) (S5 Fig). Interestingly, in the pre-gb zone the aspect ratio (max L/W) of *mac>DfosDN* macrophages was not different from control macrophages (Fig 5C-D) although the *mac>DfosDN* cells were 9% smaller in both their length and width (S5D Fig). This suggested that the gb could impose resistance on the entering macrophage, an effect which *mac>DfosDN* macrophages have trouble overcoming due to their compromised cortical actin cytoskeleton.

**Fig 5.**
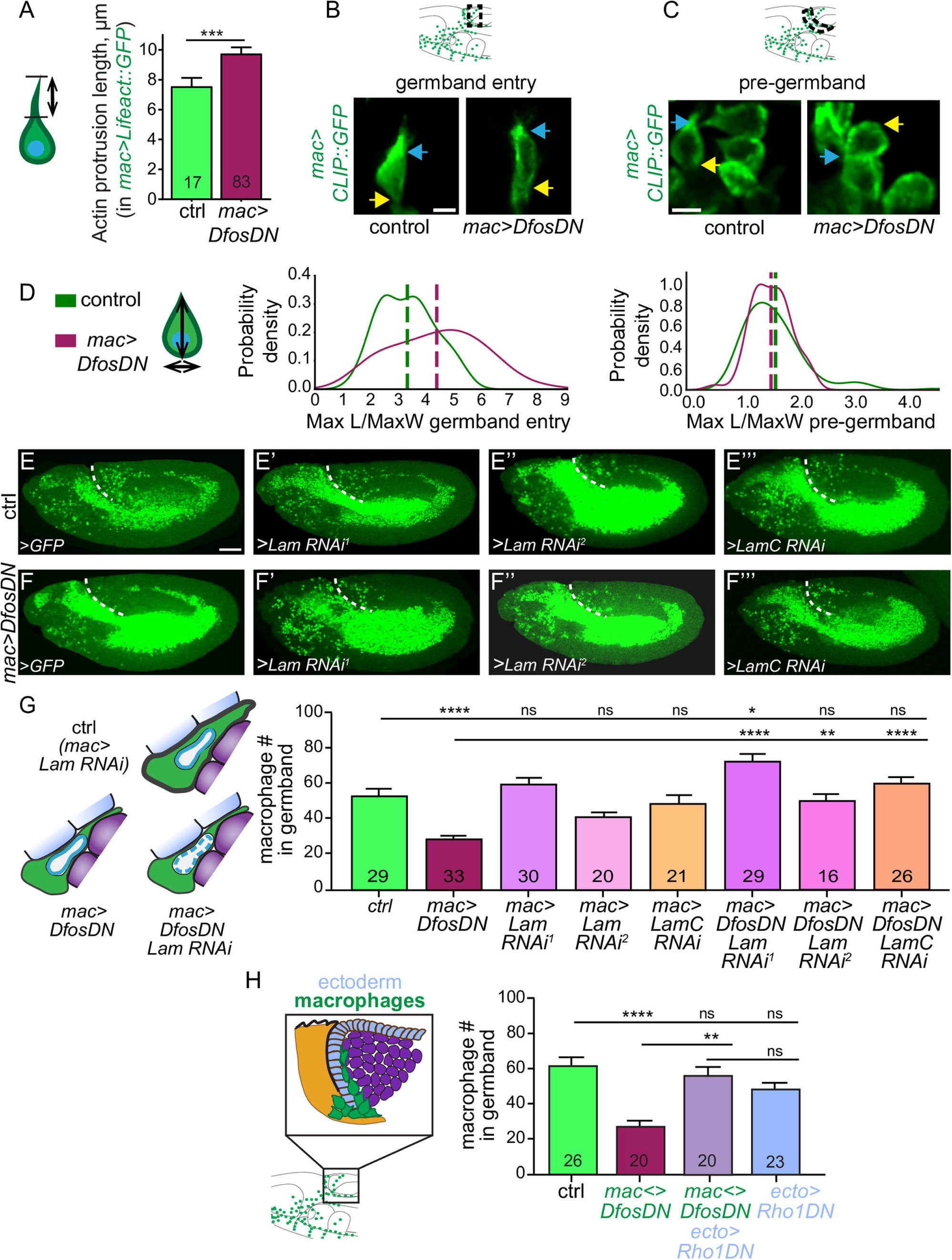
Dfos aids macrophage gb invasion against the resistance of surrounding tissues and buffers the nucleus. **(A)** Quantification from live embryos shows that the length of the F-actin protrusion of the first entering macrophage is longer in macrophages expressing DfosDN. p= 0.011. The F-actin protrusion labelled with *srpHemo-Gal4* driving *UAS-LifeAct::GFP* was measured in the direction of forward migration (see schematic). SD: 2.4, 3.7 **(B-C)** Stills from 2-photon movies of St 11 embryos showing **(B)** the first macrophages entering the gb and **(C)** macrophages in the pre-gb zone in the control and in a line expressing DfosDN in macrophages. Microtubules are labelled with *srpHemo-Gal4* driving *UAS-CLIP::GFP*. A blue arrow indicates the front and a yellow arrow indicates the rear of the macrophage. Schematics above indicate where images were acquired **(D)** Schematic at left shows macrophage measurements: vertical line for the maximum length and horizontal line for the maximum width. Histograms show the probability density distributions of the aspect ratios (maximum length over maximum width) of the first macrophage entering the gb (left) and macrophages in the pre-gb (right). Macrophages expressing *DfosDN* are more elongated the *mac>DfosDN* line. Control vs. *DfosDN* aspect ratios at gb entry p=0.0011, in pre-gb p=0.53. SD: in gb 1.0, 1.6; in pre-gb 0.5, 0.5. Confocal images of St 12 embryos expressing RNAi against Lamin or LaminC in macrophages in **(E-E’’’)** the control, or **(F-F’’’)** in embryos also expressing DfosDN in macrophages. *srpHemo-GAL4* used as drover. *Lam RNAi^1^*: GD45636; *Lam RNAi^2^*KK107419. *Lam C RNAi:* TRiP JF01406 **(G)** Macrophage RNAi knockdown of Lamins which can increase nuclear deformability did not affect macrophages numbers in the gb in the control. In embryos in which macrophages expressed DfosDN, Lamin knockdown rescues their reduced numbers in the gb. Control vs. *DfosDN* p<0.0001. Control vs. *Lam RNAi^1^* p>0.99, vs. *Lam RNAi^2^* p=0.83, vs. *LamC RNAi* p>0.99. Control vs. *DfosDN, Lam RNAi^1^* p=0.024, vs. *DfosDN, Lam RNAi^2^* p>0.99, vs. *DfosDN, LamC RNAi* p>0.99. *DfosDN* vs. *DfosDN, Lam RNAi^1^* p<0.0001, vs. *DfosDN, Lam RNAi^2^* p=0.0049, vs. *DfosDN, LamC RNAi* p<0.0001. SD: 22, 10, 19, 11, 21, 23, 16, 20. **(H)** Expressing *DfosDN* in macrophages reduces their number in the gb. Concomitantly reducing tissue tension in the ectoderm (light blue in schematic) through Rho1DN substantially rescues invasion. *srpHemo-QF QUAS* control (*mac<>)* governed macrophage expression and *e22C-GAL4* ectodermal (*ecto>*). Control vs. *mac<>DfosDN* p<0.0001 (56% reduction), vs. *mac<>DfosDN; ecto>Rho1DN* p>0.99, vs. *ecto>Rho1DN* p=0.11. *mac<>DfosDN* vs. *mac<>DfosDN; ecto>Rho1DN* p<0.0001, vs. *ecto>Rho1DN* p=0.0044. *mac<>DfosDN; ecto>Rh1oDN* vs. *ecto>Rho1DN* p>0.99. SD: 23, 16, 21, 18. Macrophages are labeled in **B-C** by *srp-Gal4* driving *UAS-CLIP::GFP,* and in **E-F’”** by *srpHemo-Gal4 UAS-mCherry-nls*. ***p<0.005, **p<0.01, *p<0.05. Unpaired t-test was used for **A,** one way ANOVA with Tukey post hoc for **G-H**. The number shown within the column corresponds to measurements in **A,** and analysed embryos in **G-H.** Scale bar 5μm in **B-C**, and 50μm in **E-F**’’’.

### Dfos promotes advancement of macrophages against the resistance of the surrounding tissues and buffers the nucleus

We therefore examined how the properties of the gb tissues and macrophages interact during invasion. We first investigated if the macrophage nucleus impedes normal invasion by varying levels of the two *Drosophila* Lamin genes, Lam and LamC, both equally related to the vertebrate lamins A and B1 (Muñoz-Alarcón et al., 2007) and both shown to affect nuclear stiffness and deformability (Wintner et al., 2020; Zwerger et al., 2013). Over-expressing Lam (S5E Fig) or knocking down either of these Lamins in macrophages through RNAi (Perkins et al., 2015) did not change macrophage numbers in the gb of wild type embryos (Fig 5E-E’’’, G), suggesting that the properties of the macrophage nucleus are not a rate limiting parameter during normal tissue invasion into the narrow path between the ectoderm and mesoderm. This result also argues that Lamins’ capacity to alter gene expression is not normally important for invasion (Andrés & González, 2009). However in *mac>DfosDN* macrophages, knockdown of these Lamins was able to rescue the gb invasion defect (Fig 5E-G), supporting the conclusion that the properties of the nucleus affect invasion in the absence of the higher levels of cortical actin Dfos normally induces. To directly test if reducing the tension of surrounding tissues can counteract the absence of Dfos, we expressed Rho1DN in the ectoderm with the *e22C-GAL4* driver while expressing *QUAS-DfosDN* in macrophages with the GAL4-independent Q-system driver we had constructed, *srpHemo-QF2* (Gyoergy et al., 2018). Rho1 through ROCK is a key regulator of Myosin activity, epithelial tension and tissue stiffness (Warner & Longmore, 2009; Zhou et al., 2009); Myosin II is essential for actin contractility (Heer & Martin, 2017) and tension in the *Drosophila* gb ectoderm (Ratheesh et al., 2018). Indeed, we found that this reduction of ectodermal tension substantially rescued DfosDN expressing macrophage numbers in the gb (Fig 5H). Taken together our results argue that Dfos aids *Drosophila* macrophages in withstanding the resisting force of surrounding cells against the nucleus during invasion into tissues.

## Discussion

We identify the ability to tune the state of the cortical actin cytoskeleton as a key capacity for immune cells migrating into and within tissue barriers *in vivo*. We find that macrophages upregulate a program governed by the transcription factor Dfos to enable this. Dfos in *Drosophila* is known to regulate the movement during dorsal or wound closure of epithelial sheets (Brock et al., 2012; Lesch et al., 2010; Riesgo-Escovar & Hafen, 1997; Zeitlinger et al., 1997) as well as the development of epithelial tumors and their dissemination (Külshammer et al., 2015; Uhlirova & Bohmann, 2006; Külshammer & Uhlirova, 2013; Benhra et al., 2018). Here we define a different role, namely that Dfos enables a stream of individual immune cells to efficiently push their way into tissues, a process which is aided rather than hampered by the presence of the ECM (Sánchez-Sánchez et al., 2017; Valoskova et al., 2019). This function appears to be specifically required for invasion as we observe no defects in *DfosDN* macrophages’ migratory speed in open environments. *DfosDN* macrophages display decreased actin at the cell circumference and an elongated shape within in the confinement of the germband, suggesting a defect in the stiffness of the cortex. Strikingly, only in the presence of *DfosDN* does the state of the nucleus become relevant, with reductions in Lamins shown to underlie nuclear stiffness (Wintner et al., 2020) enhancing the ability of macrophages to invade. These findings along with the ability of a softened ectoderm to substantially rescue the *DfosDN* macrophages’ germband invasion defect lead us to propose the model (Fig 6) that Dfos permits efficient initial translocation of the macrophage body under ectodermal reactive load by forming a stiff cortical actin shell that counteracts surrounding tissue resistance and protects the nucleus from undergoing high levels of mechanical stress during tissue entry.

**Fig. 6.**
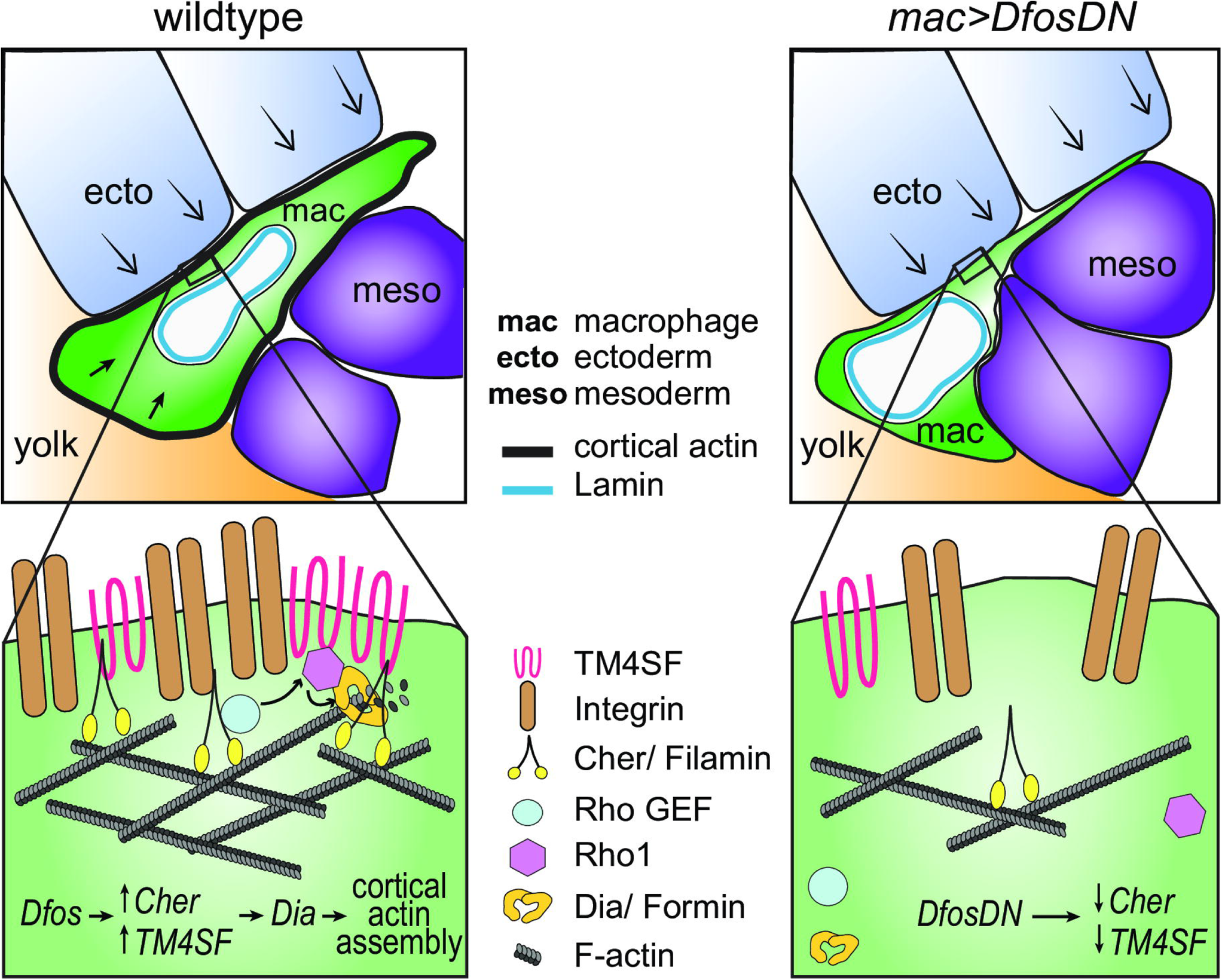
Model: Dfos increases actin assembly and crosslinking through the tetraspanin TM4SF and the Filamin Cheerio to counter surrounding tissue resistance. We propose a model for how Dfos tunes the cortical actin properties of *Drosophila* embryonic macrophages to aid their infiltration against the resistance of the surrounding germband tissue. Dfos leads to an increase of the tetraspanin TM4SF and the Filamin Cheerio (Cher). The binding of TM4SF and Filamin to different partners (see Figure S6) forms a network at the cell surface of Integrin, actin and upstream signaling molecules; this results in the recruitment of Rho GEFs and activation of Rho1 GTPase and the formin Diaphanous, which can stimulate further actin polymerization at the cortex. Thereby, F-actin is assembled into a more crosslinked and dense network aiding the macrophage in moving its cell body into the ecto-meso interface. The presence of Lamin around the nuclear membrane does not affect this process in the wildtype since the dense cross-linked cortical actin network helps macrophages withstand the load of the surrounding tissues. However, in the DfosDN-expressing macrophages, the loss of Cher and TM4SF lead to reduced cross-linked actin levels at the cell cortex making the stiffness of the nucleus the rate limiting step for macrophage infiltration of the gb tissue.

### A molecular program for tissue invasion that strengthens cortical actin

Crucial mediators of this process are two actin regulators, the filamin Cher, known to be a Dfos target in epithelia, and the previously uncharacterized membrane scaffold tetraspanin TM4SF. We show that both require Dfos for higher mRNA levels in macrophages and present correlative evidence that these classes of genes are also upregulated by vertebrate c-fos. Each of these Dfos targets is required for macrophage invasion; over-expression of TM4SF in macrophages can rescue the *DfosDN* tissue invasion phenotype. We propose that these targets act together to strengthen the actin cytoskeleton for tissue invasion. Higher Filamin levels cross-link actin filaments into resilient and stiffer networks maintaining cell integrity during mechanical stress (Goldmann et al., 1997; Tseng et al., 2004; Fujita et al., 2012). This aids the distribution of forces from focal adhesions across the entire migrating cell body, since Filamins can bind directly to Integrin, and even more strongly under strain (Ehrlicher et al., 2011; Glogauer et al., 1998; Kumar et al., 2019; Razinia et al., 2012). Tetraspanins, self-associating multipass transmembrane proteins, also can bind Integrin, forming microdomains of adhesion molecules, receptors and their intracellular signaling complexes, including Rho GTPases (Zimmerman, et al., 2016; Termini & Gillette, 2017; Yáñez-Mó et al., 2009; Berditchevski & Odintsova, 1999).

Filamins similarly bind receptors, regulators of actin assembly, Rho GTPases and the Rho GEF Trio (Popowicz et al., 2006; Stossel et al., 2001; Vadlamudi et al., 2002; Ohta et al., 1999; Bellanger et al., 2000). We observe reduced cortical levels of F-actin, active Rho1 GTPase, and the actin polymerizing formin Diaphanous in the absence of either Dfos, the Filamin Cheerio or the Tetraspanin TM4SF. Thus our data supports the hypothesis that these Dfos targets enhance the cortical recruitment and activation of the formin Dia to stimulate actin polymerization at least in part through activating the Rho1 GTPase (Fig 6, S6 Fig) (Rousso et al., 2013; Seth et al., 2006; Großhans et al., 2005; Williams et al., 2007; Delaguillaumie et al., 2002). Dfos’ upregulation of both targets could thus lead to a supra-network in which ECM-anchored FAs connect to a strong cross-linked cortical actin lattice, allowing Myosin contraction to be converted into cellular advancement despite resistance from the flanking ectoderm.

We demonstrate that the actin nucleating formin Dia is important for *Drosophila* macrophage invasion and capable of rescuing the defects in the *DfosDN* mutant. Unlike the formin Ena which mediates chemotaxis (Davidson et al., 2019), Dia is not required for general *Drosophila* macrophage migration, and instead allows macrophages to recoil away from one another (Davis et al., 2015). Dia could be required for macrophages specifically when they face resistance from their surroundings and need to increase their cortical tension. Modeling indicates that Dia1’s regulation of cortical tension requires an optimal combination of actin cross linking and intermediate actin filament length (Chugh et al., 2017). *Drosophila* Dia is a more processive nucleator than Ena (Bilancia et al., 2014 and thus could create the intermediate length actin filaments that enable higher levels of macrophage cortical tension and strain stiffening (Kasza et al., 2010) on all sides of the cell during their invasion.

Our findings thus demonstrate that there are commonalities in the molecular mechanisms by which *Drosophila* cells invade into either confluent tissues or the ECM. Dfos’s upregulation of the Filamin Cheerio is also required in tumor cells and aneuploid epithelial cells to enhance ECM breaching (Külshammer & Uhlirova, 2013; Benhra et al., 2018). Both cell types displayed enhanced levels of cortical filamentous actin, which in the tumors is concomitant with Dia upregulation (Külshammer & Uhlirova, 2013). In the oocyte, Filamin is required for follicle cell intercalation and border cells display higher levels of Filamin and F-actin to maintain cellular integrity during migration between nurse cells (Sokol & Cooley, 2003; Somogyi & Rørth, 2004). The mediator of these increased F-actin levels, MAL-D, can be activated by Dia (Somogyi & Rørth, 2004). Thus while MMPs may be specific to ECM crossing, a denser and more cross linked actin cortex due to increased levels of the filamin Cheerio and activity of the formin Dia could be a common feature of *Drosophila* cells moving through the resistance of either ECM or surrounding tissues. Determining if such shifts in cell surface actin properties underlie some vertebrate cancer cells’ capacity to metastasize even in the presence of MMP inhibitors is an interesting area of inquiry (Butcher et al 2009; Kessenbrock et al 2010).

### Implications for vertebrate immune cell migration

Our work also suggests a new perspective on the migration of some vertebrate immune cells. We find that altering lamin levels does not normally affect *Drosophila* macrophage tissue invasion. This contrasts with results showing that nuclear deformability from lower lamin levels underlies the migration of some immune cell types through narrow constrictions engineered from rigid materials (Davidson et al., 2014; Thiam et al, 2016). However, negotiation of such extremely challenging *in vitro* environments can lead to DNA damage (Raab et al., 2016) and higher nuclear flexibility caused by lower lamin levels is associated with increased cell death (Harada et al., 2014). A robust cell surface actin layer would allow long-lived cells or those not easily replenished to protect their genome as they move through resistant yet deformable environments. Embryonic *Drosophila* and vertebrate tissue resident macrophages migrate into tissues during development, survive into the adult, and serve as founders of proliferative hematopoetic niches (Holz et al., 2003; Makhijani et al., 2011; Bosch et al., 2019; Ginhoux and Guilliams, 2016; Theret et al 2019; Guilliams et al, 2020). Tissue resident memory T cells migrate in response to infection in mature animals, are long-lived and not easily renewed from the blood (Szabo et al., 2019). Thus the importance of nuclear mechanics for migration in challenging *in vivo* environments should be explored for a broader range of immune cells as well as the utilization of cortical actin as a strategy for genomic protection.

## Materials and Methods

### Fly strains and genetics

Flies were raised on standard food bought from IMBA (Vienna, Austria) containing agar, cornmeal, and molasses with the addition of 1.5% Nipagin. Adults were placed in cages in a fly room or a Percival DR36VL incubator maintained at 25°C and 65% humidity or a Sanyo MIR-153 incubator at 29°C within the humidity controlled 25°C fly room; embryos were collected on standard plates prepared in house from apple juice, sugar, agar and Nipagin supplemented with yeast from Lesaffre (Marcq, France) on the plate surface. Fly crosses and embryo collections for RNA interference experiments (7 hour collection) as well as live imaging (6 hour collection) were conducted at 29°C to optimize expression under GAL4 driver control (Duffy, 2002). All fly lines utilized are listed below.

### Fly stocks

*srpHemo-GAL4 (mac>)* was provided by K. Brückner (UCSF, USA)(Brückner et al., 2004). *Oregon R (control), P{CaryP}attP2 (control), P{CaryP}attP40 (control), kay^2^ (Dfos^2^), (UAS-Fra)2 (Dfos), UAS-Rho1.N19 Rho1DN), UAS-fbz (DfosDN), UAS-kayak* RNAi *(Dfos RNAi)* TRiP HMS00254 and TRiP JF02804, *UAS-dia RNAi* TRiP HM05027, *UAS-LamC RNAi* TRiP JF01406 and TRiP HMS00308, *e22c-GAL4 (ecto>)*, *Resille::GFP*, *UAS-GFP.nls, UAS-mCherry.nls, UAS-CD8::GFP* lines were obtained from the Bloomington Stock Center (Indiana, USA). *kay^1^* (*Dfos^1^)* line was provided by O. Schuldiner (WIS, Israel). *UAS-dia.deltaDad.EGFP (diaCA)* and *srpHemo-GAL4 UAS-CLIP::GFP (mac>CLIP::GFP)* lines were provided by B. Stramer (KCL, UK). *UAS-cher.FLAG (cher)* line was provided by M. Uhlirova (CECAD, Germany). *w[1118] (control), UAS-her* RNAi KK107451, *UAS-TM4SF* RNAi KK102206, *UAS-Lam RNAi^1^* GD45636, *UAS-Lam RNAi^2^* KK107419 lines were obtained from the Vienna *Drosophila* Resource Center (Austria).

### Extended genotypes

Here we list the lines used in each Fig; we state first the name from FlyBase; in parentheses the name used in the Fig panels is provided.

#### Fig 1 and S1 Fig

Fig 1D: *Oregon R*. Fig 1E-G: *srpHemo-GAL4, UAS-GFP (control)*. S1A, F Fig: *srpHemo-Gal4, srpHemo-H2A::3xmCherry/P{CaryP}attP2 (control)*. Fig 1H: *srpHemo-GAL4, UAS-GFP; kay^1^ (Dfos^1^)*. Fig 1I-L and S1B, G Fig: *srpHemo-GAL4, UAS-GFP::nls/+ (control 1)*. Fig 1H, 1J, 1N: *srpHemo-GAL4, UAS-GFP/+; kay^1^ (Dfos^1^)*. Fig 1K, 1N and S1B: *srpHemo-GAL4, UAS-GFP::nls/+; kay^2^ (Dfos^2^)*. Fig 1L, 1N: *srpHemo-GAL4, UAS-GFP::nls/(UAS-Fra)2*; *kay^2^ (Dfos^2^;mac>Dfos)*. Fig 1O, 1Q: *10XUAS-IVS-myr::GFP/+; srpHemo-Gal4, srpHemo*-*H2A:*:3x*mCherry/+ (control 2 and control)*. Fig 1P-Q: *UAS-DfosDN/+; srpHemo-Gal4, srpHemo*-*H2A:*:3x*mCherry/+ (mac>DfosDN)*. S1C, F Fig: *srpHemo-Gal4, srpHemo*-*H2A:*:3x*mCherry/ UAS GFP::nls* (ctrl). *srpHemo-Gal4, srpHemo*-*H2A:*:3x*mCherry/UAS-fbz (mac>DfosDN)*. S1D Fig: *srpHemo-Gal4, srpHemo*-*H2A:*:3x*mCherry /+* (ctrl). *srpHemo-Gal4, srpHemo*-*H2A:*:3x*mCherry/UAS-DfosDN (mac>DfosDN)*. Fig 1R and S1E, G, I Fig: *UAS-GFP; srpHemo-Gal4, srpHemo-H2A::3xmCherry (ctrl). UAS-Dfos RNAi HMS00254/srpHemo-Gal4, srpHemo-H2A::3xmCherry (mac>DfosRNAi^1^). UAS-Dfos RNAi JF02804/srpHemo-Gal4, srpHemo-H2A::3xmCherry (mac>DfosRNAi^2^)*. S1H Fig: *srpHemo-GAL4, UAS-GFP::nls/+* or /(*UAS-Fra)2 (mac>Dfos)*. S1I Fig: *UAS-GFP; UAS-Dfos RNAi HMS00254/srp-GAL4, srpHemo-Gal4, srpHemo-H2A::3xmCherry (mac>DfosRNAi^1^+ GFP). UAS-GFP; UAS-Dfos RNAi JF02804/srpHemo-Gal4, srpHemo-H2A::3xmCherry (mac>DfosRNAi^2^+ GFP)*.

#### Fig 2 and S2 Fig

Fig 2A, 2C-I and S2A-B, E Fig: *srpHemo-Gal4, srpHemo*-*H2A:*:3x*mCherry/+ (control)*. Fig 2D: *srpHemo-Gal4, srpHemo*-*H2A:*:3x*mCherry/+* (3 movies) and *Resille::GFP/+; srpHemo-Gal4, srpHemo*-*H2A:*:3x*mCherry/+* (4 movies, control) and *Resille::GFP/+; srpHemo-Gal4, srpHemo*-*H2A:*:3x*mCherry/+* (3 movies) and *Resille::GFP/+; srpHemo-Gal4, srpHemo*-*H2A:*:3x*mCherry/UAS-DfosDN (*4 movies, *DfosDN)* Fig 2A, 2C-I and S2A-B, E Fig: *srpHemo-Gal4, srpHemo*-*H2A:*:3x*mCherry/UAS-fbz (mac>DfosDN)*. S2C-D Fig: *srpHemo-GAL4, UAS-GFP.nls/+ (control)*. S2C-D Fig: *srpHemo-GAL4, UAS-GFP.nls/+; kay^2^ (Dfos^2^)*.

#### Fig 3 and S3 Fig

Fig 3C, G and S3D Fig: *UAS-Dicer2;; srpHemo-Gal4, srpHemo*-*H2A:*:3x*mCherry/w^1118^ (control)*. Fig 3D, 3G and S3D Fig: *UAS-Dicer2; UAS-TM4SF RNAi* KK10220/*+; srpHemo-Gal4, srpHemo*-*H2A:*:3x*mCherry/+ (mac>TM4SF RNAi)*. Fig 3E, G and S3D Fig: *UAS-Dicer2; UAS-cher RNAi* KK107451*/+; srpHemo-Gal4, srpHemo*-*H2A:*:3x*mCherry/+ (mac>cher RNAi)*. Fig 3F-G: *UAS-Dicer2; UAS-cher RNAi* KK107451/*UAS-TM4SF RNAi* KK102206; *srpHemo-Gal4, srpHemo*-*H2A:*:3x*mCherry/+ (mac>TM4SF RNAi, cher RNAi)*. Fig 3H, L: *srpHemo-GAL4, UAS-mCherry::nls/UAS-mCD8::GFP (control)*. Fig 3I, L: *srpHemo-GAL4, UAS-mCherry::nls/UAS-mCD8::GFP; UAS-fbz/+ (mac>DfosDN)*. Fig 3J, L: *srpHemo-GAL4,UAS-mCherry::nls/UAS-cheerio::FLAG; UAS-fbz/+ (mac>DfosDN, cher)*. Fig 3K-L: *srpHemo-GAL4,UAS-mCherry.nls/UAS-TM4SF; UAS-fbz/+ (mac>DfosDN, TM4SF)*. Fig 3L: *srpHemo-GAL4, UAS-mCherry::nls/ UAS-TM4SF (mac>TM4SF)*. Fig 3L: *srpHemo-GAL4, UAS-mCherry::nls/UAS-cher (mac>cher)*. S3A-C Fig: *srpHemo-Gal4, srpHemo*-3x*mCherry/+ (control)*. S3A-C Fig: *srpHemo-Gal4, srpHemo*-3x*mCherry/UAS-fbz (mac>DfosDN)*.

#### Fig 4 and S4 Fig

Fig 4A: *srpHemo-3xmCherry; kay^1^ (Dfos^1^) and srpHemo-3xmCherry; +*. Fig 4B, D and S4A Fig: *srpHemo-Gal4, srpHemo*-*moe::*3x*mCherry/+;UAS-mCD8::GFP/+ (Control)*. Fig 4C-D and S4A Fig: *srpHemo-Gal4, srpHemo*-*moe::*3x*mCherry/UAS-fbz (mac>DfosDN)*. S4A Fig: *w^118^*. Fig 4E-F: *srpHemo-Gal4, srpHemo*-*moe::*3x*mCherry/w^118^ (Control)*. Fig 4E, G: *srpHemo-Gal4, srpHemo*-*moe::*3x*mCherry/UAS-cher RNAi* KK107451 *(mac>cher RNAi)*. Fig 4E, H: *srpHemo-Gal4, srpHemo*-*moe::*3x*mCherry/UAS-TM4SF RNAi* KK102206 *(mac>TM4SF RNAi)*. Fig 4I-J: *srpHemo-GAL4, UAS-mCherry.nls/UAS-mCD8::GFP (control)*. Fig 4I’, J: *srpHemo-GAL4, UAS-mCherry.nls/UAS-DiaΔDad::EGFP; UAS-fbz/+ (mac>DfosDN, diaCA*). Fig 4J: *srpHemo-GAL4, UAS-mCherry.nls/UAS-mCD8::GFP; UAS-fbz/+ (mac>DfosDN)*. Fig 4J: *srpHemo-GAL4, UAS-mCherry.nls/ UAS-DiaΔDad::EGFP (mac>diaCA)*. S4B Fig: #1: *UAS-GFPnls; srpHemo-Gal4, srpHemo-H2A::3xmCherry*. #2: *UAS-GFPnls*/*srpHemo-Gal4, srpHemo-H2A::3xmCherry*; *Dfos^1^*. #3: *UAS-GFPnls*/ *srpHemo-Gal4, srpHemo-H2A::3xmCherry*; *Dfos^2^*. #4: *UAS-DiaΔDad::EGFP*/*srpHemo-Gal4, srpHemo-H2A::3xmCherry*; *Dfos^1^*. #5: *UAS-Dia⊗Dad::EGFP*/*srpHemo-Gal4, srpHemo-H2A::3xmCherry*; *Dfos^2^*. Fig 4K-L and S4C-D Fig: *UAS-Dicer2;; srpHemo-Gal4, srpHemo*-*H2A:*:3x*mCherry/P{CaryP}attP40 (control)*. Fig 4K’, L and S4C-D Fig: *UAS-Dicer2;+; srpHemo-Gal4, srpHemo*-*H2A:*:3x*mCherry/ UAS-dia RNAi* HM05027 *(mac>dia RNAi^1^)*. Fig 4L and S4C-D Fig: *UAS-Dicer2;+; srpHemo-Gal4, srpHemo*-*H2A:*:3x*mCherry/UAS-dia RNAi* HMS00308 *(mac>dia RNAi^2^)*. Fig 4M-N and S4E Fig: *(control) UAS-diaRBD::GFP/+; UAS-nlacz/ srpHemo-Gal4, 10XUAS-IVS-myr::tdTomato. (mac>Rho1DN) UAS-diaRBD::GFP/+; UAS-Rho1N.19)/srpHemo-Gal4, 10XUAS-IVS-myr::tdTomato. (mac>DfosDN) UAS-diaRBD::GFP/+; UAS-fbz/srpHemo-Gal4, 10XUAS-IVS-myr::tdTomato. (mac>cher RNAi) UAS-diaRBD::GFP/+; UAS-her RNAi KK107451/srpHemo-Gal4, 10XUAS-IVS-myr::tdTomato. (mac>TM4SF RNAi) UAS-diaRBD::GFP/+; UAS-TM4SF RNAi KK102206/srpHemo-Gal4, 10XUAS-IVS-myr::tdTomato*. Fig 4O-P and S4F Fig: *(control) UAS-dia::GFP/+; srpHemo-Gal4, 10XUAS-IVS-myr::tdTomato/UAS-nlacZ. (mac>Rho1DN) UAS-dia::GFP/+; srpHemo-Gal4, 10XUAS-IVS-myr::tdTomato/UAS-Rho1N.19. (mac>DfosDN) UAS-dia::GFP/UAS-fbz; srpHemo-Gal4, 10XUAS-IVS-myr::tdTomato/+*. *(mac>cher RNAi) UAS-dia::GFP/UAS-cher RNAi KK107451; srpHemo-Gal4, 10XUAS-IVS-myr::tdTomato/+. (mac>TM4SF RNAi) UAS-dia-GFP/UAS-TM4SF RNAi KK102206; srpHemo-Gal4, 10XUAS-IVS-myr::tdTomato/+*.

#### Fig 5 and S5 Fig

Fig 5A and S5A Fig: *srpHemo-Gal4 UAS-LifeActGFP UAS-RedStinger/ srpHemo-Gal4 UAS-LifeActGFP, UAS-RedStinger* control*; srpHemo-Gal4 UAS-LifeActGFP UAS-RedStinger/ srpHemo-Gal4 UAS-LifeActGFP UAS-RedStinger; UAS-DfosDN/UAS-DfosDN*. Fig 5B-D and S5B-D Fig: *srpHemo-Gal4, UAS-CLIP::GFP, UAS-RedStinger (control)*. Fig 5B-D and S5B-D Fig: *srpHemo-Gal4, UAS-CLIP::GFP, UAS-RedStinger; UAS-fbz (mac>DfosDN)*. Fig 5E, G: *srpHemo-GAL4, UAS-mCherry.nls/UAS-mCD8::GFP (control)*. Fig 5E’-E’’, 5G: *srpHemo-GAL4, UAS-mCherry.nls/UAS-Lamin RNAi* GD45636, KK107419 *(mac>Lam RNAi^1^* and *mac>Lam RNAi^2^, respectively)*. Fig 5E’’’, G: *srpHemo-GAL4, UAS-mCherry.nls/UAS-LaminC RNAi* TRIP JF01406 *(mac>LamC RNAi)*. Fig 5F-G: *srpHemo-GAL4, UAS-mCherry.nls/UAS-mCD8::GFP; UAS-fbz/+ (mac>DfosDN)*. Fig 5F’,F’’, G: *srpHemo-GAL4, UAS-mCherry.nls/UAS-Lam RNAi* (*Lam RNAi^1^=*GD45636, *Lam RNAi^2^*=KK107419)*; UAS-fbz/+ (mac>DfosDN, Lam RNAi^1^* and *mac>DfosDN, Lam RNAi^2^)*. Fig 5F’’’, G: *srpHemo-GAL4, UAS-mCherry.nls/UAS-LaminC RNAi* TRIP JF01406*; UAS-fbz/+ (mac>DfosDN, LamC RNAi)*. Fig 5H: *e22CGal4,srpHemo*-*H2A:*:3x*mCherry/+ (control)*. Fig 5H: *srpQF/ srpHemo*-*H2A:*:3x*mCherry; QUAS-fbz/UAS-Rho1.N12 (mac<>DfosDN)*. Fig 5H: *e22CGal4, srpHemo*-*H2A:*:3x*mCherry/srpQF; +/ UAS-Rho1.N12 (ecto>Rho1DN)*. Fig 5H: *srpQF/ e22C-Gal4, srpHemo*-*H2A:*:3x*mCherry; UAS-Rho1N12/QUAS-fbz (mac<>DfosDN, ecto>rho1DN)*. S5E Fig: *+;UAS-GFP::nls, srpHemo-GAL4 (control). +;UAS-GFP::Lamin, srpHemo-GAL4*.

### Cloning and generation of QUAS-DfosDN line

The fragment was amplified from genomic DNA of the published *UAS-fbz (UAS-Dfos DN)* line (Eresh, Riese, Jackson, Bohmann, & Bienz, 1997) using primers encompassing a 5’ consensus translation initiation sequence followed by the bZIP fragment and containing BglII and XhoI restriction sites: 5’-GAAGATCTATTGGGAATTCAACATGACCCCG-3’ and 5’-CCCTCGAGTCAGGTGACCACGCTCAGCAT-3’. The resulting fragment was cloned into the pQUASt vector, a gift from Christopher Potter (Addgene plasmid # 104880). The final construct was sequenced and injected into the attP2 landing site by BestGene (Chino Hills, CA, USA).

### Cloning and generation of UAS-TM4SF line

The TM4SF open reading frame was amplified from the DGRC GH07902 cDNA clone (#3260, Fbcl0121651), using primers acagcgGAATTCATGGCATTGCCGAAGAAAAT and acagcgTCTAGATTAAAAGCTAATCGTCTGTCATT. The PCR product and the pUASt-aTTB vector (DGRC plasmid #1419) were digested with EcoRI and XbaI, and ligated. After sequencing, the construct was injected into the landing site line, (*y^1^ M{vas-int.Dm}ZH-2A w\**; *M{3xP3-RFP.attP}ZH-51D*, BL 24483), to produce second chromosome inserts. All male survivors were crossed to *w*; *Sp/CyO*; *PrDr/TM3Ser* virgins. Transformants were recognized by eye color and crossed again to *w*; *Sp/CyO*; *PrDr/TM3Ser* virgins to get rid of the X chromosomal integrase.

### Embryo staging

Laterally oriented embryos with complete germband (gb) extension and the presence of stomadeal invagination were staged based on gb retraction from the anterior as a percentage of total embryo length. Embryos with no gb retraction were classified as Stage 11, 30% retraction early Stage 12, 60% retraction Stage 12, and 70% Stage 13. Imaged embryos are shown throughout paper in a lateral orientation with anterior to the left and dorsal up.

### In situ hybridization and immunofluorescence

Embryos were dechorionated by 5 min treatment with 50% Chlorox bleach. After extensive washing with water, embryos were fixed with 3.7% formaldehyde/heptane for 20 min followed by methanol devitellinization for *in situ* hybridization and visualization of 3xmCherry or tdTomato. The *Dfos* cDNA clone SD04477 was obtained from the DGRC. T7 or T3 polymerase-synthesized digoxigenin-labelled anti-sense probe preparation and *in situ* hybridization was performed using standard methods (Lehmann & Tautz, 1994). Images were taken with a Nikon-Eclipse Wide field microscope with a 20X 0.5 NA DIC water Immersion Objective. Embryos were mounted after immunolabeling in Vectashield Mounting Medium (Vector Labs, Burlingame, USA) and imaged with a Zeiss Inverted LSM700 and LSM800 Confocal Microscope using a Plain-Apochromat 20X/0.8 Air Objective or a Plain-Apochromat 63X/1.4 Oil Objective as required.

| Antibody | Source animal | Dilution | Provided by |
| --- | --- | --- | --- |
| Anti-Dfos | Rabbit | 1:50 | J. Zeitlinger (Stowers Institute, USA) |
| Anti-GFP | Chicken | 1:500 | Abcam (ab13970) |
| Anti-mCherry | Goat | 1:200 | Invitrogen (M11217) |

### Dfos antibody

The Dfos rabbit polyclonal antibody was produced for the lab of Julia Zeitlinger. It was raised by Genescript (Piscataway, NJ, USA) against the C-terminal end of *Drosophila* Kayak found in all isoforms and was purified against an N terminally His tagged antigen corresponding to aa 73 to 595 of Kay isoform A. The internal Genescript order number is 163185-30, and in the Zeitlinger lab is referred to as anti-kay/fos Ab.

### Western

Cages were prefed on fresh yeast plates for two days. Late stage 11/ early Stage 12 embryos were hand picked using a Leica M205 fluorescent microscope on ice cold apple juice plates. They were transferred to RIPA buffer (50mM Tris, 150mM NaCl, 1% NP-40, 1mM EDTA, 0.5% Na-Deoxycholate, 0.1% SDS) with a Halt Protease/Phosphatase inhibitor cocktail (ThermoFisher, #78440) and lysed. After a 30 min incubation on ice, they were centrifuged 15 min at 4°C at 15,000 g. 10 µg of the cleared lysate were separated by SDS-PAGE using 4%-15% Mini-PROTEAN TGX Precast Protein gels (Bio-Rad, #4561085) and blotted onto a Amersham Protran Premium Western blotting nitrocellulose membrane (Sigma, #GE10600003). The nitrocellulose membrane was blocked with Pierce Clear Milk blocking buffer (ThermoFisher, #37587) and incubated in blocking buffer with anti-mCherry (Novus Biologicals, #NBP1-96752) at 1:1000, and anti-Profilin (DSHB, #chi 1J) at 1:50 antibodies over night at 4°C. The membrane was washed three times for 10 minutes with 1x PBS and incubated with Goat Anti-Mouse IgG (H+L)-HRP Conjugate (BioRad, #172-1011). Chemiluminescence was induced by incubation with SuperSignal West Femto Maximum Sensitivity Substrate (ThermoFisher, #34096) and recorded with a ChemieDoc MP (BioRad) molecular imager. Densitometric quantification of bands was done with ImageJ.

### Time-Lapse Imaging

Embryos were dechorionated in 50% bleach for 5 min, washed with water, and mounted in halocarbon oil 27 (Sigma) on a 24×50mm high precision coverslip (Marienfeld Laboratory Glassware, No. 1.5H) between two bridges (~0.5 cm high) of coverslips glued on top of each other, or mounted in halocarbon oil 27 (Sigma) between a 18×18mm coverslip (Marienfeld Laboratory Glassware, No. 1.5H) and an oxygen permeable membrane (YSI). The embryo was imaged on an upright multiphoton microscope (TrimScope, LaVision) equipped with a W Plan-Apochromat 40X/1.4 oil immersion objective (Olympus). GFP and mCherry were imaged at 860 nm and 1100 nm excitation wavelengths, respectively, using a Ti-Sapphire femtosecond laser system (Coherent Chameleon Ultra) combined with optical parametric oscillator technology (Coherent Chameleon Compact OPO). Excitation intensity profiles were adjusted to tissue penetration depth and Z-sectioning for imaging was set at 1µm for tracking. For long-term imaging, movies were acquired for 60 - 150 minutes with a frame rate of 25-45 seconds. A temperature control unit set to 29°C was utilized for all genotypes except *kay^2^* for which the setting was 25°C.

### Image Analysis

**Macrophage cell counts:** Autofluorescence of the embryo revealed the position of the germband (gb) for staging of fixed samples. Embryos with 40% (±5%) gb retraction (Stage 12) were analysed for macrophage numbers in the pre-gb, within the germband, along the ventral nerve cord (vnc) and in the whole embryo. For the *kay RNAi*.embryos with 70% gb retraction (Stage 13) were used for vnc counts. The pre-gb zone was defined based on embryo and yolk autofluorescence as an area on the yolk sac underneath the amnioserosa with borders defined posteriorly by the gb ectoderm and anteriorly by the head. Macrophages were visualized using confocal microscopy with a Z-stack step size of 2 µm and macrophage numbers within the gb or the segments of the vnc were calculated in individual slices (and then aggregated) using the Cell Counter plugin in FIJI. Total macrophage numbers were obtained using Imaris (Bitplane) by detecting all the macrophage nuclei as spots.

### Macrophage Tracking, Speed, Persistence. Mode of Migration and Macrophage gb crossing Analysis

Embryos with macrophage nuclei labelled with *srpHemo-H2A::3XmCherry* and the surrounding tissues with *Resille::GFP,* or with only macrophages labelled by *srpHemo-H2A::3XmCherry, or srpHemo>GFP.nls* were imaged and 250×250×40µm^3^ 3D-stacks were typically acquired with ~0.2×0.2×1µm^3^ voxel size every 39-41 seconds for ~2 hours. For imaging macrophages on vnc frames were acquired at every 40-43 seconds for 30 min after macrophages started spreading into abdominal segment 2 (see Fig 2G). Multiphoton microscopy images were initially processed with ImSpector software (LaVision Bio Tec) to compile channels, and exported files were further processed using Imaris software (Bitplane) for 3D visualization.

Each movie was rotated and aligned along the embryonic AP axis for tracking analysis. For analysis of migration in the pre-gb and gb in the control and *kay^2^* mutant, embryos were synchronized using the onset of germ and retraction. For vnc migration analysis, macrophages were tracked for 30 minutes from when macrophages started moving into the second abdominal segment. Only macrophages migrating along the inner edge of the vnc were analyzed.

Gb crossing time was calculated from when the macrophages align in front of the gb ectoderm in a characteristic arc, until the first macrophage had transitioned its nucleus inside the ecto-meso-interphase. To see the gb edge and yolk in movies of *srpHemo-3xH2A::mCherry*, either *Resille::GFP* labelling the outlines of all cells, or the auto-fluorescence of the yolk was used.

For analysis of gb migration in the *DfosDN* vs control macrophages, macrophages were tracked from when the first macrophage appeared between the ectoderm and the yolk sac until gb retraction started, typically 60 minutes. In the head and pre-gb, macrophage nuclei were extracted using the spot detection function, and tracks generated in 3D over time. The pre-gb and gb were defined as for macrophage counts described above. The mean position of the tracks in X- and Y restrict analysis to each migratory zones.

Cell speed and persistence were calculated from nuclei positions using custom Python scripts as described elsewhere (Smutny et al., 2017). Briefly, instantaneous velocities from single cell trajectories were averaged to obtain a mean instantaneous velocity value over the course of measurement. The directional persistence of a trajectory was calculated as the mean cosine of an angle between subsequent instantaneous velocities:

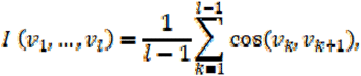

where is duration of the trajectory and are its instantaneous velocities. Only trajectories with a minimal duration of 15 timeframes were used. Calculated persistence values were averaged over all trajectories to obtain a persistence index (*I*) for the duration of measurement (with −1 being the lowest and 1 the maximum). 3-6 embryos were recorded and analyzed for each genotype, numbers of control and perturbed embryos are equal in each pairwise comparison.

### Measurement of junctional Phalloidin

The junctional intensity of F-actin (Phalloidin) was calculated using linescan analysis as previously described (Smutny et al., 2010) with the following changes. The line was ~5 µm and was always drawn in the middle slice of the Z stack (1 µm resolution) of the macrophage-macrophage junction. For every line, a Gaussian fit was applied and maximum intensities across the cell junction were then normalized against average intensities of F-actin (Phalloidin) staining in the stereotypical gb area of ~50×50µm^2^ in each embryo. Analyses were carried out using standard Fiji software. 4-5 embryos were analysed per genotype. Macrophages in the pre-gb or gb entry zones were analyzed.

### Measurement of F-actin reporters

To quantify cortical F-actin intensity in living embryos, a *srpHemo-moe::3xmCherry* reporter line (Gyoergy et al., 2018) was crossed into a background of macrophages expressing *DfosDN*, *cher RNAi*, or *TM4SF* RNAi. Embryos were collected for 5h 30min at 29°C, de-chorionated in 50% bleach for 5 min, rinsed thoroughly with water, and aligned laterally side by side under a stereomicroscope using a fluorescence lamp to check for the presence of mCherry. Aligned embryos were then mounted as described in the live imaging section above. To image Moe::3xmCherry, a Zeiss LSM800 inverted microscope was used with the following settings: Plan-APOCHROMAT 40×/1.4 Oil, DIC, WD=0.13 objective, 1.5× zoom, 1025×1025 pixel, speed 8, heating chamber set to 29°C, z-interval 1μm. Laser settings were kept constant in all experiments. Images were acquired during macrophage invasion into the gb (St 12). Pseudo-coloring was conducted for the mCherry red channel. Each pixel in the image has a color ascribed to it via the fire “Look Up Table” translating the level of intensity of the mCherry channel into a defined amount of each color. The highest intensity of the image is represented as very bright yellow and all other grey values are depicted as colors on the scale accordingly.

For quantification of Moe::3xmCherry intensity, an ROI was drawn in Fiji software around macrophages at the germband entry site in 20 z-stacks for each embryo. The area mean intensity was measured in all ROIs and the average/embryo was calculated. To normalize fluorescence intensities per batch, the average intensity/embryo of all ROIs in each sample was divided by the arithmetic mean of the average intensity/embryo of all ROIs in the control per batch. The normalized average intensities/embryo were then compared to each other using a t-test with Welch’s correction for *DfosDN* and one way-ANOVA for *cher RNAi* and *TM4SF RNAi*.

### Quantification of membrane localization of DiaRBD::GFP and Dia::GFP

Methanol fixed St 11 embryos were mounted either after staining with GFP antibody (Dia::GFP) or without staining(DiaRBD::GFP) and imaged with a Zeiss Inverted LSM800, Plain-Apochromat 63X/1.4 Oil Objective at an XY-resolution of 0.1 µm and a Z-resolution of 1 µm (~15 µm total stack). All macrophages within 40µm of the germband were analysed. For the quantification of the levels of DiaRBD or the complete Dia protein at the plasma membrane versus the cytoplasm, confocal images were processed using Fiji and MATLAB-R2017b (MathWorks). Individual focal planes were used to segment a profile corresponding to an 8 pixel wide line drawn across the single outer membrane of individual macrophages chosen such that the extracellular portion of the line extended into surrounding tissue or space and not another macrophage. The corresponding intensity profiles of the mem::Tomato and Dia::GFP or DiaRBD::GFP channels were extracted in Fiji using a custom macro and analyzed further using a custom MATLAB script. The membrane region was defined by finding the maximal value in the Tomato intensity profile and centering a 0.8 μm interval around it. The background was calculated for each GFP profile as the mean intensity in the 2 µm outside the cell, flanking the membrane region, and substracted from the entire profile. The integrated Dia::GFP or DiaRBD::GFP intensity at the membrane was calculated within the 0.8 μm interval defined above. The integrated cytoplasmic Dia::GFP or DiaRBD::GFP level was calculated as the mean intensity of 2 µm of the GFP profile inside the cell flanking the membrane region. Image analysis scripts are publicly available at https://github.com/Axmasha/Image_analysis_scripts.

### Cell aspect ratio analysis and imaging actin dynamics

Laterally oriented embryos were used to measure the maximal length and width of macrophages expressing *UAS-CLIP::GFP* under the control of *srpHemoGal4*. Briefly, 3D-stacks with 1 µm Z resolution were acquired every 35-45 seconds for approximately 1 hour. As the strength of the GAL4 expression increased over time, laser power was adjusted during acquisition to reach the best possible quality of visualization. Images acquired from mutiphoton microscopy were initially processed with ImSpector software (LaVision Bio Tec) to compile channels from the imaging data.

We started measuring from the time the cell body of the first macrophage fully appeared at the interface between the ectoderm and mesoderm and yolk sac until it had moved 30 µm along the ectoderm mesoderm interface. At each timeframe, a line was drawn in Fiji along the longest dimension of the macrophage in the direction of its front-rear polarization axis, denoted the maximal cell length, and along the orthologonal longest dimension, which was considered maximal cell width. We did not observe long CLIP::GFP protrusions, but when a small protrusion was present, it was not included in the length measurement; within this gb region the front of the first macrophage was clearly outlined with CLIP::GFP. The border between the first and second entering macrophages was drawn based on the uninterrupted intense line of CLIP::GFP at the base of the first macrophage; only cells with a clearly visible border were measured. The length to width ratio was quantified for each timeframe and a probability density function was plotted: 5 embryos were recorded for each genotype.

### Imaging the actin protrusion

Laterally oriented embryos expressing *srpHemo-Gal4 UAS-LifeAct::GFP* were used to image macrophage actin live with a 3D-stack resolution of 1µm. See above description of CLIP::GFP labeled macrophage imaging for laser power and image compilation. Laser power was also increased further in the DfosDN samples to enhance actin visualization. We measured the length of the filopodia-like protrusion of the first entering macrophage with Imaris software (Bitplane) from the time when the protrusion was inserted into the ectoderm, mesoderm and yolk sac interface until the macrophage started to translocate its cell body into that location.

### FACS sorting of macrophages

Adult flies of either *w;+;srpHemoGal4,srpHemo::3xmCherry/+* or *w;+; srpHemoGal4,srpHemo::3xmCherry* /*UASDfosDN* genotypes were placed into plastic cages closed with apple juice plates with applied yeast to enhance egg laying. Collections were performed at 29°C for 1 hour, then kept at 29°C for additional 5 hours 15 minutes to reach stage 11-early stage 12. Embryos were harvested for 2 days with 6-7 collections per day and stored meanwhile at +4°C to slow down development. Collected embryos were dissociated and the macrophages sorted as previously described (Gyoergy et al., 2018). About 1-1.5×10^5^ macrophages were sorted within 30 minutes.

### Sequencing of the macrophage transcriptome

Total RNA was isolated from FACS-sorted macrophages using Qiagen RNeasy Mini kit (Cat No. 74104). The quality and concentration of RNA was determined using Agilent 6000 Pico kit (Cat No. 5067-1513) on an Agilent 2100 Bioanalyzer: on average about 100 ng of total RNA was extracted from 1.5×10^5^ macrophages. RNA sequencing was performed by the CSF facility of Vienna Biocenter according to standard procedures (https://www.vbcf.ac.at/facilities/next-generation-sequencing/) on three replicates. Briefly, the cDNA library was synthesized using QuantSeq 3’ mRNA-seq Library Prep kit and sequenced on the Illumina HiSeq 2500 platform. The reads were mapped to the *Drosophila melanogaster* Ensembl BDGP6 reference genome with STAR (version 2.5.1b). The read counts for each gene were detected using HTSeq (version 0.5.4p3). Flybase annotation (r6.19) was used in both mapping and read counting. Counts were normalised to arbitrary units using the TMM normalization from edgeR package in R. Prior to statistical testing the data was voom transformed and then the differential expression between the sample groups was calculated with limma package in R. The functional analyses were done using the topGO and gage packages in R (Anders, Pyl, & Huber, 2015; Dobin et al., 2013). RNA sequencing data has been deposited at GEO as GSE182470.

### qRT-PCR analysis of mRNA levels in murine bones and osteosarcomas

RNA isolation and qPCR was performed from bones of wild-type C57BL/6 mice and from bones and osteosarcomas (OS) of H2-c-fosLTR as previously described with the below primers (Rüther et al., 1989).

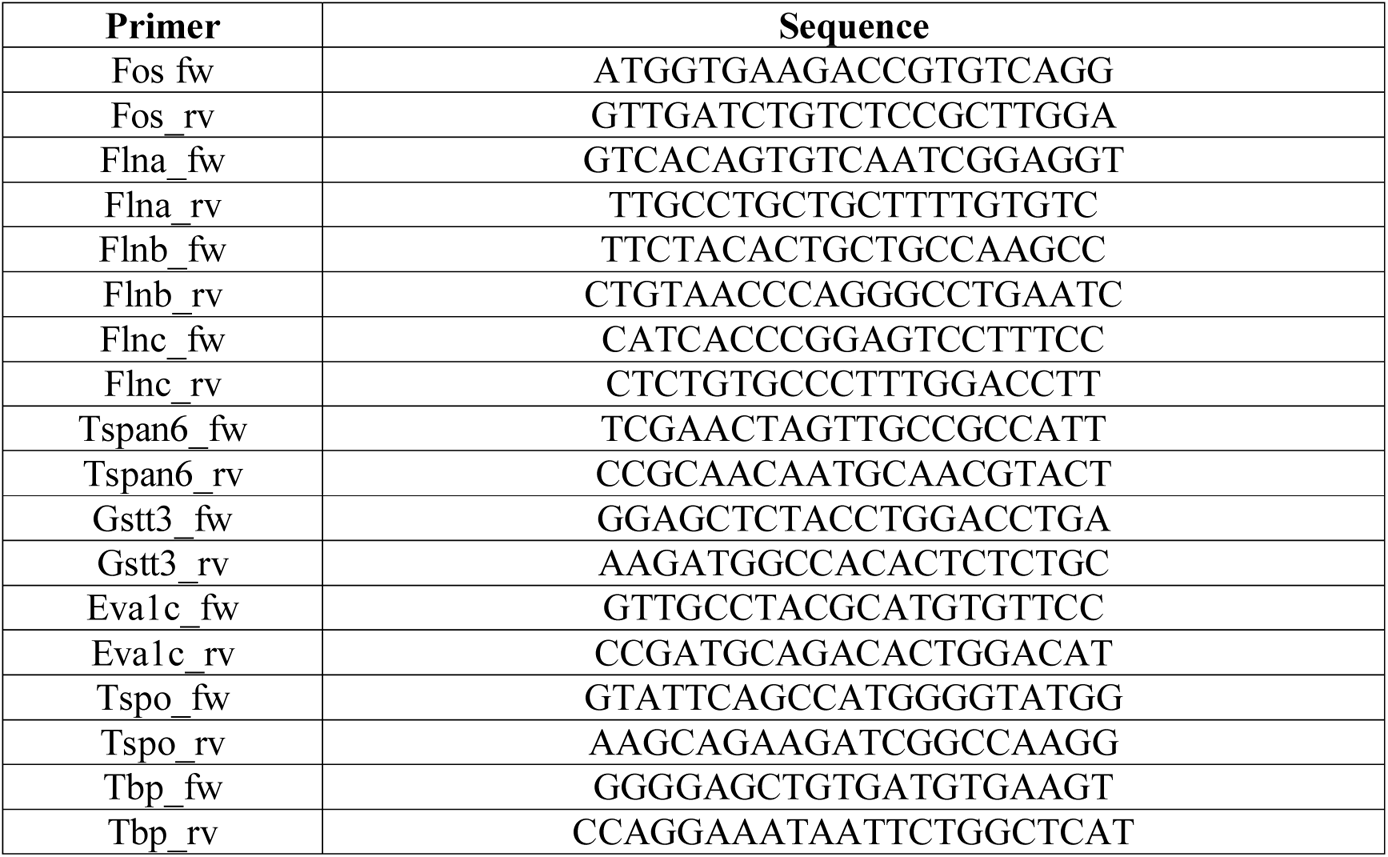

### Statistics and Repeatability

#### Mouse experiments

Data are shown as mean±SEM. One-way ANOVA followed by Tukey’s multiple comparisons post-test was applied to compare experimental groups. Statistical analysis was performed using GraphPad Prism 6.0 software. A p-value <0.05 was considered statistically significant (*p<0.05, **p<0.01, ***p<0.001, ****p<0.0001).

#### *Drosophila* experiments

Statistical tests as well as the number of embryos/cells/tracks/contacts assessed are listed in the Figure legends. All statistical analyses were performed using GraphPad PRISM or R Studio and significance was determined using a 95% confidence interval. No statistical method was used to predetermine sample size.

Representative images of Dfos antibody staining were analyzed per replicate per genotype and *in situ* hybridization are from experiments that were repeated 2 times with many embryos with reproducible results. *Dfos* mutants analysis in Fig 1 and S1 Fig are from experiments that were repeated 2-3 times. In live imaging experiments in Fig 2 and S2 Fig, 3-7 embryos for each genotype were analyzed, each embryo was recorded on a separate day. FACS sorting of macrophages from embryos was conducted in three replicates, from which RNA samples were prepared for RNA sequencing. Experiments in Fig 4 and S4 Fig were repeated at least 3 times, with representative images and plots of phalloidin immunostaining from experiments that were repeated 4 times. In the LifeAct::GFP protrusion live imaging experiment in Fig 5 and S5 Fig, 3-5 embryos were analyzed for each genotype. In CLIP::GFP live imaging experiments in Fig 5 and S5 Fig, 5-6 embryos were analyzed for each genotype for the cell aspect ratio in the germband zone, and 2 embryos in the pre-germband zone and for tracking of the front vs rear speed. Each embryo was recorded on a separate day. The Lamin over expression in S5 Fig and the Lamin knockdown rescue experiments in Fig 5G were repeated at least 3 times. The gb rescue experiment in Fig 5H was repeated at least 4 times.

## Acknowledgements

We thank the following for their contributions: The *Drosophila* Genomics Resource Center supported by NIH grant 2P40OD010949-10A1 for plasmids, K. Brueckner. B. Stramer, M. Uhlirova, O. Schuldiner, the Bloomington *Drosophila* Stock Center supported by NIH grant P40OD018537 and the Vienna *Drosophila* Resource Center for fly stocks, FlyBase (Thurmond et al., 2019) for essential genomic information, and the BDGP *in situ* database for data (Tomancak et al., 2002, 2007). For antibodies, we thank the Developmental Studies Hybridoma Bank, which was created by the Eunice Kennedy Shriver National Institute of Child Health and Human Development of the NIH, and is maintained at the University of Iowa, as well as J. Zeitlinger for her generous gift of Dfos antibody. We thank the Vienna BioCenter Core Facilities for RNA sequencing and analysis and the Life Scientific Service Units at IST Austria for technical support and assistance with microscopy and FACS analysis. We thank C. P. Heisenberg, P. Martin, M. Sixt and Siekhaus group members for discussions and T. Hurd, A. Ratheesh and P. Rangan for comments on the manuscript.

## Supplementary Information Titles/Legends

**S1 Data. Data used to plot all graphs and to perform statistical analyses**

**S2 Data. RNA sequencing data**

**S1 Fig. Dfos does not affect the total number of macrophages, or their number in the pre-gb zone and along the vnc**

**(A)** Dfos protein (red) is detected with an antibody in macrophages (green) in embryos from the stages as indicated.

(**B-I**) Quantification in mid St 12 embryos.

**(I) (B)** The number of macrophages (green) in the pre-gb zone (outlined by a black dotted line in the schematic on the left) showed no significant change in *Dfos^2^* mutant embryos compared to the control (p=0.37) SD: 6,7.

**(II) (C)** The total number of macrophages (see schematic at left) was not altered from that in the control embryos expressing DfosDN in macrophages (p=0.12). SD: 60, 120.

(**D-E**) The number of macrophages (green) along the vnc (outlined by black dotted line in the schematic on the left) shows no significant difference between the control and **(D)** macrophages that express DfosDN or **(E)** either of two RNAi lines against *Dfos*. (**D**) *DfosDN* p=0.88, 0.99, >0.99. *Dfos RNAi^1^* (TRiP HMS00254) p=0.21, 0.06, 0.11, 0.072, 0.033, 0.30, 0.56. *Dfos RNAi^2^* (TRiP JF02804) p=0.34, 0.15, 0.83, 0.27, 0.47, 1.0, 0.45. (D) SD: Ctrl 3, 3, 3, 0.8; *DfosDN* 6, 3, 0.7. (E) SD: Ctrl 6, 3, 3, 3, 2, 0.3; *Dfos RNAi^1^* 6, 3, 3, 3, 2, 2, 0.3; *Dfos RNAi^2^* 6, 2, 3, 2, 3, 1, 0.4.

(**F, G**) Macrophage numbers in the pre-gb (see schematic at left) are increased compared to the control for lines expressing (**F**) DfosDN or (**G**) one of two different *UAS-Dfos RNAi* constructs in macrophages under *srpHemo-GAL4* control. (F) p=.04, SD: 19, 29. (G) *Dfos RNAi^1^* p<0.0009, *Dfos RNAi^2^* p<0.0001. SD: 12, 9, 14.

**(A) (H)** Macrophage numbers in the germband are not significantly altered compared to the control upon overexpression of Dfos in macrophages (p=0.14). SD: 22, 14.

**(I)** Macrophage numbers in the gb for lines expressing one of two different *UAS-Dfos RNAi* constructs in macrophages under *srpHemo-GAL4* control and lines which additionally express *UAS-GFP*. Control vs. *mac>Dfos RNAi^1^* (TRiP HMS00254) or Control vs. *mac>Dfos RNAi^2^* (TRiP JF02804), p<0.0001. *mac>Dfos RNAi^1^* vs. *mac>Dfos RNAi^1^ + GFP* or *mac>Dfos RNAi^2^* vs. *mac>Dfos RNAi^2^ + GFP*, p>0.99. SD: 33, 47, 34. The effect of each *Dfos RNAi* was eliminated upon simultaneous expression of another UAS construct.

Macrophages are labeled using either *srpHemo-Gal4* driving *UAS-GFP* or *srpHemo-H2A::3xmCherry*. “*mac>*” indicates *srpHemo-GAL4* driver expressing UAS constructs specifically in macrophages. Histograms show mean + SEM ***p<0.005, **p<0.01, *p<0.05. Unpaired t-test was used for statistics, except for G, I, which used One-Way ANOVA. The number of embryos analysed for that genotype is shown within each column in the graphs. In D n=6 embryos for the control and n=9 for Dfos DN. In E n=9 embryos for control, 15 and 11 for *Dfos RNAis*. Scale bar in A: 10 µm.

**S2 Fig. Dfos facilitates macrophage motility during initial invasion into the tissue**

**(A)** Quantification reveals that the directional persistence of macrophages expressing DfosDN (0.58) is unchanged (0.56) in the pre-gb area (p=0.66) but decreased during gb entry (0.65) (0.72), p=0.038 and along the vnc (0.54) compared to the control (0.61), p=0.00026. Left schematic shows pre-gb area in yellow, gb entry outlined in solid line. Boxed area in right schematic shows analyzed area of vnc.

**(B)** Movie stills showing wild type and *Dfos^2^* macrophages entering the gb (outlined by the dashed line). Time in min shown in the top right corner of each image.

**(C)** Quantification of macrophage speed shows a significant reduction in the speed of *Dfos^2^* macrophages in the pre-gb zone and at gb entry, but none in the head. Regions analysed indicated in left schematic. Speed in head: control=2.59 μm/min, *Dfos* =2.68 μm/min, p=0.40; speed in pre-gb=3.38 μm/min, *Dfos* =2.47 μm/min, p=2.38e; speed in gb entry: control=2.35 μm/min, *Dfos* =1.62 μm/min, p= 0.0003.

Macrophages are labeled using *srpHemo-H2A::3xmCherry*. Histograms show mean + SEM ***p<0.005, **p<0.01, *p<0.05. Unpaired t-test was used for statistics. The number of analyzed macrophages for each genotype shown within each graph column. Tracks were obtained from movies of 3 embryos each for control and *mac>DfosDN* for pre gb entry in **A**, 4 each for gb entry in **A**, 3 each for the vnc in **A**, 4 each of control and 4 *Dfos^2^* embryos for head and pre-gb in **C**, and 3 embryos each for gb entry in **C**. Scale bars: 10 µm.

**S3 Fig. Dfos regulates macrophage germband invasion through actin cytoskeleton associated proteins**

**(A-C)** Comparative mRNA expression levels as determined from RNA sequencing analysis of FACS sorted wild type macrophages and those expressing DfosDN, n=3 biological replicates. (**A**-**B**) Genes down-regulated in macrophages expressing DfosDN are shown, separated into those with **(A)** strong and **(B)** moderate expression in wild type macrophages.

**(A) (C)** Expression levels of *Drosophila* formin family genes are unchanged. Fold enrichment is normalized. p-values: *Dhc36C* 0.02, *CG14204* 0.03, *CG42402* 0.04, *CR43767* 0.046, *TM4SF* 0.03, *CG42260* 0.0011, *cher* 0.046, *GstT4* 0.018, *Xrp1* 0.0011, *Tspo* 0.046, CG31337 0.046. *Frl*, *DAAM*, *dia*, *capu* all >0.99.

**(D-E)** Quantification of the macrophage numbers in **(D)** the pre-gb zone and **(E)** along the vnc from embryos expressing RNAi against *cher* (*KK 107451*), or *TM4SF* in macrophages (*KK 102206*) driven by *srpHemo-Gal4* shows no significant alteration. The number in the column in **(D)** corresponds to the number of embryos analyzed. Control vs. *cher RNAi* p=0.33. Control vs. *TM4SF RNAi* p=0.05. Control vs. *cher/TM4SF RNAi* p=0.67. (D) SD: 20, 20, 19, 13.

For **(E)** n=13 embryos for control and n=15 for each *cher RNAi* and *TM4SF RNAi*. Control vs. *cher RNAi* p=0.97 for T1, p=0.33 for T2, p=0.88 for T3. Control vs. *TM4SF RNAi* p=0.52 for T1, p=0.76 for T2, p=0.35 for T3. SD: ctrl 6.5, 5.4, 0.6; *cher RNAi* 5.0, 3.3, 0.8; *TM4SF RNAi* 4.4, 4.9, 1.9.

**(F-I)** q-PCR analysis of mRNA extracted from the bones of mice that are wild type, transgenic (tg) for MHC c-fos, viral 3’UTR, and those in which c-fos transgenesis has led to an osteosarcoma (OS). Analysis of mRNA expression shows that **(F)** higher *Fos* levels in osteosarcomas correlate with hgher levels of (**G**) the glutathione S transferase *Gstt3*, and **(H)** the slit receptor *Eva1c*. but not **(I)** *Tspo*. Bone and osteosarcoma RNA isolated from the same transgenic mouse, n=4 mice per group, age 5 to 6 months. p-values= 0.86, 0.0028, 0.0013 in **(F)**, 0.79, 0.0001, 0.0003 in **(G)**, 1.0, 0.054, 0.049 in **(H)**, 0.37, 0.33, 0.040 in **(I)**. SD: 0.7, 0.6, 2.6 in **(F)**; 0.2, 0.3, 1.1 in **(G)**; 0.4, 0.2, 1.5 in **(H)**; 0.1, 0.2, 0.2 in **(I)**.

Histograms show mean + SEM ***p<0.005, **p<0.01, *p<0.05.

Unpaired t-test or one-way ANOVA with Tukey post hoc were used for statistics of quantifications. Significance is based on adjusted p values.

**S4 Fig. Dia does not affect macrophage numbers in the pre-gb zone and along the vnc**

**(A)** Three Western blots probed with an mCherry antibody of St 11 embryo extracts from *srpHemo-moe::3xmCherry* expressing either *CD8::GFP* (ctrl) or *DfosDN* in macrophages. Left Western blot also contains *w-*lane. **A’**) Quantitation of the Western blots. We observed no significant change in the expression of the Moe protein reporter when Dfos function is inhibited.

**(B)** Expressing Dia-CA in macrophages in *Dfos^1^* or *Dfos^2^* embryos completely rescued the macrophage germband invasion defect. p-values: Control vs. *Dfos^1^* or vs. *Dfos^2^* p=0.0004 or p=0.0055 respectively; Control vs. *Dfos^1^ mac>DiaCA* or vs. *Dfos^2^ mac>DiaCA* p >0.999; *Dfos^1^* vs. *Dfos^1^ mac>DiaCA* p=0.0005; *Dfos^2^* vs. *Dfos^2^ mac>DiaCA* p=0.035. SD: 20, 23, 18, 19, 7.8.

(**C-D**) There was no significant change in the number of macrophages in (**C**) the pre-gb zone or (**D**) along the vnc in embryos expressing either of two different RNAi lines against *dia* expressed in macrophages. Pre-gb: Control vs. *dia RNAi^1^* p=0.54, Control vs. *dia RNAi^2^* p=0.77. vnc: Control vs. *dia RNAi^1^* p=0.99, Control vs. *dia RNAi^2^* p=0.95. *RNAi^1^* =TRiP HMS05027, *RNAi^2^* = TRiP HMS00308. (C) SD: 9, 12, 13. (D) SD: Ctrl 5.2, 6.4, 2.5, 0.4; *dia RNAi^1^* 5.6, 6.8, 1.7, 0.2; *dia RNAi^2^* 5.1, 4.9, 2.1, 0.6.

(**E-F**) Two further examples of line profiles used for the determination of the membrane-to-cytoplasmic ratios in Fig 4N, P. Line intensity profiles of (E) DiaRBD::GFP or (F) Dia::GFP (green) and membrane myr::Tomato (magenta) across the edge of macrophages expressing either lacZ (Control), RhoDN, DfosDN, *cher RNAi*, or *TM4SF RNAi* as shown in the schematic in E. Line length ~ 8 µm. Blue lines indicate mean GFP intensity on the membrane and in cytoplasm.

Histograms show mean + SEM ***p<0.005, **p<0.01, *p<0.05. One way ANOVA with Tukey post hoc was used for statistics of quantification. The number in each column corresponds to the number of analysed embryos. “mac>” indicates *srpHemo-GAL4* driver expressing *UAS* constructs in macrophages. Macrophages are labeled using *srpHemo-H2A::3xmCherry*.

**S5 Fig. Dfos controls cell shape in macrophages**

**(A)** Representative image showing actin protrusions of the first macrophage entering the germband in the control and in lines expressing DfosDN in macrophages. Actin was visualized by *srpHemo-Gal4* (“mac>”) driving *UAS-LifeActGFP*. White stars indicate the tip of each actin protrusion. Scale bar 5μm.

**(B)** Microtubules are labeled with *srpHemo-Gal4* driving *UAS-CLIP::GFP*. Spatially matched stills of the first macrophage expressing DfosDN and control extending protrusions into the gb slightly before entering with the body of the cell. As DfosDN macrophages have a delay in entry, the stills from the DfosDN movie are from a later developmental time point than the control.

**(C)** Quantification of macrophage maximum length and maximum width shows that DfosDN expressing macrophages are 23% longer and 12% thinner than wild type macrophages inside the gb (indicated in schematic above by dashed box). Control vs. *DfosDN* maximum length p=0.0005, SD: 3.4, 5.7; control vs. *DfosDN* maximum width p=0.0025, SD: 1.3, 1.0.

**(D)** Quantification of the maximum length and maximum width of macrophages in the pre-gb zone (indicated in schematic by dashed box) shows that macrophages expressing DfosDN are 9% shorter and 9% thinner than wild type macrophages. Control vs. *DfosDN* maximum length p=0.0095, SD: 2.2, 2.0; control vs. *DfosDN* maximum width p=0.005, SD: 2.3, 1.9.

**(E)** Overexpression of *UAS-Lam* in macrophages through *srpHemo-Gal4* (*mac>*) causes no change in their number in the gb compared to the control. p=0.65, SD: 15, 18.

Histograms show mean + SEM ***p<0.005, **p<0.01, *p<0.05. Unpaired t-test was used for statistics of quantification. The number of measurements per genotype is shown in each columns

### S6 Fig: Model of protein interactions at the macrophage cortex

Proposed interactions of proteins at the cell cortex in wildtype macrophages during gb infiltration as shown in Fig 6. Direct binding between two proteins is indicated by a line, signalling between the interaction partners is represented as an arrow. These interactions and the resulting model in Figure 6 are based on the papers at the end of this legend next to the corresponding number shown for each linkage. The Tetraspanin TM4SF can cluster adhesion receptors such as Integrins at the membrane and lead to the recruitment and activation of Rho GTPases. Rho GTPases can bind and activate the formin Dia leading to F-actin polymerization. In addition, Integrin can bind filamins (Cher), which can bind to and thereby recruit RhoGEF to the membrane. Rho GEFs can in turn bind to and activate Rho GTPases. References for listed interactions: **1, Tetraspanins-Integrin)** Berditchevski & Odintsova, 1999; Termini & Gillette, 2017**. 2, Tetraspanins-Rho GTPases)** Shigeta et al., 2003; Sit & Manser, 2011; Hong et al., 2012; Tejera et al., 2013**. 3, Tetraspanins-Filamins)** Brzozowski et al., 2018; Perez-Hernandez et al., 2013. **4, Integrin-Filamins)** Razinia et al., 2012; Ehrlicher et al., 2011. **5, Rho GTPases-Formins)** Kuhn & Geyer, 2014; Rose et al., 2005. **6 & 8, Rho GEF-Rho GTPases)** Vetter & Wittinghofer, 2001 **7, Formins-Filamins)** Hu et al., 2014; Lian et al., 2016**. 9, Filamins-RhoGEFs)** Bellanger et al., 2000; Min et al., 2002.

**S1 Movie. Dfos facilitates macrophage motility during initial invasion into the germband tissue**

Movies corresponding to stills shown in **Fig 2A**. Macrophages (green) labelled using *srpHemo-H2A::3xmCherry* are imaged while entering the gb in control embryos (left) and embryos in which macrophages express a dominant negative (DN) version of Dfos (DfosDN) (right). Time in minutes is indicated in the upper right corner. Scale bar: 10 µm.

**S2 Movie. Dfos does not affect macrophage migration along the vnc**

Movies corresponding to stills shown in **Fig 2G**. Macrophages (green) labelled by *srpHemo-Gal4* driving *UAS-GFP::nls* are imaged during their migration along the segments of the vnc in control embryos (left) and embryos in which DfosDN is expressed in macrophages (right). Time in minutes is indicated in the upper right corner. Scale bar: 10 µm.

**S3 Movie. Macrophages in *Dfos^2^* mutants invade germband more slowly**

Movies corresponding to stills shown in **S2B Fig**. Macrophages (green) labelled by *srpHemo-Gal4* driving *UAS-GFP::nls* are imaged while entering the gb in control embryos (left) and *Dfos^2^* mutant embryos (right). Time is indicated in minutes. Scale bar: 10 µm.

**S4 Movie. DfosDN expressing macrophages make long actin protrusions during germband entry**

Movies corresponding to stills shown in **S5A Fig**. F-actin in macrophages (green) labelled with *srpHemo-Gal4* driving *UAS-LifeAct::GFP* is imaged during gb entry in control embryos (left) and embryos with macrophages expressing DfosDN (right). Note the extended protrusion of the DfosDN expressing macrophages. Time is indicated in minutes. Scale bar: 10 µm.

**S5 Movie. Dfos controls cell shape in macrophages**

Movies corresponding to stills shown in **Fig 5B-C** and **S5B Fig**. Microtubules of macrophages are labelled with *srpHemo-Gal4* driving *UAS-CLIP::GFP*. They are imaged during gb entry in control embryos (left) and embryos with macrophages expressing DfosDN (right). Note the extended shape of the DfosDN expressing macrophages. Time is indicated in minutes. Scale bar: 10 µm.

## References

1. Anders, S., Pyl, P. T., & Huber, W. (2015). HTSeq-A Python framework to work with high-throughput sequencing data. Bioinformatics, 31(2), 166–169. https://doi.org/10.1093/bioinformatics/btu638

2. Abreu-Blanco, M.T., Verboon, J.M., & Parkhurst S.M. (2014) Coordination of Rho Family GTPase activities to orchestrate cytoskeleton responses during wound repair. Current Biology, 24(2), 144–155. https://doi.org/10.1016/j.cub.2013.11.048

3. Andrés, V., & González, J. M. (2009). Role of A-type lamins in signaling, transcription, and chromatin organization. Journal of Cell Biology, 187(7), 945–957. https://doi.org/10.1083/jcb.200904124

4. Bellanger, J. M., Astier, C., Sardet, C., Ohta, Y., Stossel, T. P., & Debant, A. (2000). The Rac1- and RhoG-specific GEF domain of trio targets filamin to remodel cytoskeletal actin. Nature Cell Biology, 2(12), 888–892. https://doi.org/10.1038/35046533

5. Benhra, N., Barrio, L., Muzzopappa, M., & Milán, M. (2018). Chromosomal Instability Induces Cellular Invasion in Epithelial Tissues. Developmental Cell, 47(2), 161–174.e4. https://doi.org/10.1016/j.devcel.2018.08.021

6. Berditchevski, F., & Odintsova, E. (1999). Characterization of integrin-tetraspanin adhesion complexes: Role of tetraspanins in integrin signaling. Journal of Cell Biology, 146(2), 477–492. https://doi.org/10.1083/jcb.146.2.477

7. Bershadsky, A. D., Balaban, N. Q., & Geiger, B. (2003). Adhesion-Dependent Cell Mechanosensitivity. Annual Review of Cell and Developmental Biology, 19(1), 677–695. https://doi.org/10.1146/annurev.cellbio.19.111301.153011

8. Bilancia, C. G., Winkelman, J. D., Tsygankov, D., Nowotarski, S. H., Sees, J. A., Comber, K., … Peifer, M. (2014). Enabled negatively regulates diaphanous-driven actin dynamics in vitro and in vivo. Developmental Cell, 28(4), 394–408. https://doi.org/10.1016/j.devcel.2014.01.015

9. Bosch, P.S., Makhijani, K., Herboso, L., Gold, K.S., Baginsky, R., Woodcock, K.J., Alexander, B., Kukar, K., Corcoran, S., Jacobs, T., Ouyang, D., Wong, C., Ramond, E.J.V., Rhiner, C., Moreno, E., Lemaitre, B., Geissmann, F., & Brueckner, K. (2019). Adult *Drosophila* lack hematopoiesis but rely on a blood cell reservoir at the respiratory epithelia to relay infection signals to surrounding tissues. Developmental Cell, 51(6)787–803. https://doi.org/10.1016/j.devcel.2019.10.017

10. Brock, A. R., Wang, Y., Berger, S., Renkawitz-Pohl, R., Han, V. C., Wu, Y., & Galko, M. J. (2012). Transcriptional regulation of profilin during wound closure in Drosophila larvae. Journal of Cell Science, 125(23), 5667–5676. https://doi.org/10.1242/jcs.107490

11. Brückner, K., Kockel, L., Duchek, P., Luque, C. M., Rørth, P., & Perrimon, N. (2004). The PDGF/VEGF receptor controls blood cell survival in Drosophila. Developmental Cell, 7(1), 73–84. https://doi.org/10.1016/j.devcel.2004.06.007

12. Brzozowski, J. S., Bond, D. R., Jankowski, H., Goldie, B. J., Burchell, R., Naudin, C., … Weidenhofer, J. (2018). Extracellular vesicles with altered tetraspanin CD9 and CD151 levels confer increased prostate cell motility and invasion. Scientific Reports, 8(1). https://doi.org/10.1038/s41598-018-27180-z

13. Butcher, D.T., Alliston, T. & Weaver, V.M.(2009). A tense situation: forcing tumour progression. Nature Reviews Cancer, 9(2), 108–22. https://doi.org/10.11038/nrc2544

14. Cho, N. K., Keyes, L., Johnson, E., Heller, J., Ryner, L., Karim, F., & Krasnow, M. A. (2002). Developmental control of blood cell migration by the Drosophila VEGF pathway. Cell, 108(6). https://doi.org/10.1016/S0092-8674(02)00676-1

15. Chugh, P., Clark, A. G., Smith, M. B., Cassani, D. A. D., Dierkes, K., Ragab, A., … Paluch, E. K. (2017). Actin cortex architecture regulates cell surface tension. Nature Cell Biology, 19(6), 689–697. https://doi.org/10.1038/ncb3525

16. Danuser, G., Allard, J., & Mogilner, A. (2013). Mathematical Modeling of Eukaryotic Cell Migration: Insights Beyond Experiments. Annual Review of Cell and Developmental Biology, 29(1). https://doi.org/10.1146/annurev-cellbio-101512-122308

17. Davidson, A. J., Millard, T. H., Evans, I. R., & Wood, W. (2019). Ena orchestrates remodelling within the actin cytoskeleton to drive robust Drosophila macrophage chemotaxis. Journal of Cell Science, 132(5). https://doi.org/10.1242/jcs.224618

18. Davidson, P.M., Denais, C., Bakshi, M., & Lammerding, J. (2014). Nuclear deformability constitutes a rate-limiting step during cell migration in 3-D environments. Cellular and Molecular Bioengineering, 7(3)293–306. https://doi.org/10.1007/s12195-014-0342-y

19. Davis, J. R., Luchici, A., Miodownik, M., Stramer, B. M., Davis, J. R., Luchici, A., … Stramer, B. M. (2015). Inter-Cellular Forces Orchestrate Contact Inhibition of Locomotion Article Inter-Cellular Forces Orchestrate Contact Inhibition of Locomotion. Cell, 161(2), 361–373. https://doi.org/10.1016/j.cell.2015.02.015

20. Delaguillaumie, A., Lagaudrière-Gesbert, C., Popoff, M. R., & Conjeaud H., H. (2002). Rho GTPase link cytoskeletal rearrangements and activation processes induced via the tetraspanin CD82 in T lymphocytes. Journal of Cell Science, 115(2).

21. Deluca, T. F., Cui, J., Jung, J. Y., St. Gabriel, K. C., & Wall, D. P. (2012). Roundup 2.0: Enabling comparative genomics for over 1800 genomes. Bioinformatics, 28(5). https://doi.org/10.1093/bioinformatics/bts006

22. Dobin, A., Davis, C. A., Schlesinger, F., Drenkow, J., Zaleski, C., Jha, S., … Gingeras, T. R. (2013). STAR: Ultrafast universal RNA-seq aligner. Bioinformatics, 29(1), 15–21. https://doi.org/10.1093/bioinformatics/bts635

23. Duffy, J. B. (2002). GAL4 system in Drosophila: A fly geneticist’s Swiss army knife. Genesis, 34(1–2), 1–15. https://doi.org/10.1002/gene.10150

24. Edwards, K. A., Demsky, M., Montague, R. A., Weymouth, N., & Kiehart, D. P. (1997). GFP-moesin illuminates actin cytoskeleton dynamics in living tissue and demonstrates cell shape changes during morphogenesis in Drosophila. Developmental Biology, 191(1). https://doi.org/10.1006/dbio.1997.8707

25. Ehrlicher, A. J., Nakamura, F., Hartwig, J. H., Weitz, D. A., & Stossel, T. P. (2011). Mechanical strain in actin networks regulates FilGAP and integrin binding to filamin A. Nature, 478(7368), 260–263. https://doi.org/10.1038/nature10430

26. Eresh, S., Riese, J., Jackson, D. B., Bohmann, D., & Bienz, M. (1997). A CREB-binding site as a target for decapentaplegic signalling during Drosophila endoderm induction. EMBO Journal, 16(8), 2014–2022. https://doi.org/10.1093/emboj/16.8.2014

27. Evans, I. R., & Wood, W. (2011). Drosophila embryonic hemocytes. Current Biology, 21(5), R173–R174. https://doi.org/10.1016/j.cub.2011.01.061

28. Franck, Z., Gary, R., & Bretscher, A. (1993). Moesin, like ezrin, colocalizes with actin in the cortical cytoskeleton in cultured cells, but its expression is more variable. Journal of Cell Science, 105(1).

29. Fujita, M., Mitsuhashi, H., Isogai, S., Nakata, T., Kawakami, A., Nonaka, I., … Kudo, A. (2012). Filamin C plays an essential role in the maintenance of the structural integrity of cardiac and skeletal muscles, revealed by the medaka mutant zacro. Developmental Biology, 361(1), 79–89. https://doi.org/10.1016/j.ydbio.2011.10.008

30. García-Echeverría, C. Methionine-containing zipper peptides. Lett Pept Sci 4, 135–140 (1997). https://doi.org/10.1007/BF02443525

31. Ginhoux, F. & Guilliams, M. (2016). Tissue-resident macrophage ontogeny and homeostasis. Immunity 44(3), 439–449. https://doi.org/10.1016/j.immuni.2016.02.024

32. Glogauer, M., Arora, P., Chou, D., Janmey, P. A., Downey, G. P., & McCulloch, C. A. G. (1998). The role of actin-binding protein 280 in integrin-dependent mechanoprotection. Journal of Biological Chemistry, 273(3), 1689–1698. https://doi.org/10.1074/jbc.273.3.1689

33. Glover, J.N.M. and Harrison, S.C. (1005). Crystal structure of the heterodimeric bZIP transcritpion factor c-Fos-c-Jun bound to DNA. Nature, 373(6511):257–261. https://doi.org/10.1038/373257a0

34. Goldmann, W. H., Tempel, M., Sprenger, I., Isenberg, G., & Ezzell, R. M. (1997). Viscoelasticity of actin-gelsolin networks in the presence of filamin. European Journal of Biochemistry, 246(2), 373–379. https://doi.org/10.1111/j.1432-1033.1997.00373.x

35. Gonzalez-Gaitan, M. & Peifer, M. (2009) Exploring the roles of Diaphanous and Enabled activity in shaping the balance between filopodia and lamellipodia. Molecular Biology of the Cell 20(24). https://doi.org/10.1091/mbc.e09-02-0144

36. Goode, B.L. & Eck. M. (2007) Mechanism and Function of Formins in the Control of Actin Assembly. Annual Review of Biochemistry 78, 593–627. https://doi.org/10.1146/annurev.biochem.75.103004.142647

37. Greten, F.R. & Grovennikov, S.I. (2019). Inflammation and Cancer: Triggers, Mechanisms and Consequences. Immunity 51(1), 27–41. https://doi.org/10.1016/j.immuni.2019.06.025

38. Großhans, J., Wenzl, C., Herz, H. M., Bartoszewski, S., Schnorrer, F., Vogt, N., … Müller, H. A. (2005). RhoGEF2 and the formin Dia control the formation of the furrow canal by directed actin assembly during Drosophila cellularisation. Development, 132(5), 1009–1020. https://doi.org/10.1242/dev.01669

39. Guilliams, M., Thierry, G.R., Bonnardel, J., & Bajenoff, M. (2020). Establishment and Maintenance of the Macrophage Niche. Immunity 52(3), 434–451. https://doi.org/10.1016/j.immuni.2020.02.015

40. Gyoergy, A., Roblek, M., Ratheesh, A., Valoskova, K., Belyaeva, V., Wachner, S., … Siekhaus, D. E. (2018). Tools allowing independent visualization and genetic manipulation of Drosophila melanogaster macrophages and surrounding tissues. G3: Genes, Genomes, Genetics, 8(3). https://doi.org/10.1534/g3.117.300452

41. Hammonds, A. A. S., Bristow, C. C. a, Fisher, W. W., Weiszmann, R., Wu, S., Hartenstein, V., … Celniker, S. E. (2013). Spatial expression of transcription factors in Drosophila embryonic organ development. Genome Biology, 14(12), R140. https://doi.org/10.1186/gb-2013-14-12-r140

42. Harada, T., Swift, J., Irianto, J., Shin, J, Spinler, K.R., Athirasala, A., Diegmiller, R., Dingal, P.C.D.P., Ivanovska, I.L., & Discher, D.E. (2014). Nuclear lamin stiffness is a barrier to 3D migration, but softness can limit survival. J. Cell Biology, (5)669–82. https://doi.org/10.1083/jcb.201308029.

43. Hattori, A., Mizuno, T., Akamatsu, M., Hisamoto, N., Matsumoto, K. (2013). The Caenorhabditis elegans JNK signaling pathway activates expression of stress response genes by depressing the Fos/HDAC repressor complex. Plos Genetics 9(2):e1003315. https://doi.org/10.1371/journal.pgen.1003315

44. Heer, N. C., & Martin, A. C. (2017). Tension, contraction and tissue morphogenesis. Development (Cambridge), 144(23)4249–4260. https://doi.org/10.1242/dev.151282

45. Holz, A., Bossinger, B., Strasser, T., Janning, W., & Klapper, R. (2003). The two origins of hemocytes in *Drosophila*. Development, 130(20), 4955–62. https://doi.org/10.1242/dev.007202

46. Homem, C.C., Peifer, M. (2008) Diaphanous regulates myosin and adherens junctions to control cell contractility and protrusive behavior during morphogenesis. Development, 135(6)1005–1018. https://doi.org/10.1242/dev.151282

47. Hong, I. K., Jeoung, D. Il, Ha, K. S., Kim, Y. M., & Lee, H. (2012). Tetraspanin CD151 stimulates adhesion-dependent activation of Ras, Rac, and Cdc42 by facilitating molecular association between β1 integrins and small GTPases. Journal of Biological Chemistry, 287(38), 32027–32039. https://doi.org/10.1074/jbc.M111.314443

48. Hu, J., Lu, J., Lian, G., Ferland, R. J., Dettenhofer, M., & Sheen, V. L. (2014). Formin 1 and filamin B physically interact to coordinate chondrocyte proliferation and differentiation in the growth plate. Human Molecular Genetics, 23(17), 4663–4673. https://doi.org/10.1093/hmg/ddu186

49. Kasza, K. E., Broedersz, C. P., Koenderink, G. H., Lin, Y. C., Messner, W., Millman, E. A., … Weitz, D. A. (2010). Actin filament length tunes elasticity of flexibly cross-linked actin networks. Biophysical Journal, 99(4), 1091–1100. https://doi.org/10.1016/j.bpj.2010.06.025

50. Kessenbrock, K., Plaks, V., & Werb, Z. (2010). Matrix metalloproteinases: regulators of the tumor microenvironment. Cell 141(1), 52–67. https://doi.org/10.1016/j.cell.2010.03.015

51. Kühn, S., & Geyer, M. (2014). Formins as effector proteins of rho GTPases. Small GTPases, 5(JUNE). https://doi.org/10.4161/sgtp.29513

52. Külshammer, E., Mundorf, J., Kilinc, M., Frommolt, P., Wagle, P., & Uhlirova, M. (2015). Interplay among Drosophila transcription factors Ets21c, Fos and Ftz-F1 drives JNK-mediated tumor malignancy. DMM Disease Models and Mechanisms, 8(10), 1279–1293. https://doi.org/10.1242/dmm.020719

53. Külshammer, E., & Uhlirova, M. (2013). The actin cross-linker Filamin/Cheerio mediates tumor malignancy downstream of JNK signaling. Journal of Cell Science, 126(4), 927–938. https://doi.org/10.1242/jcs.114462

54. Kumar, A., Shutova, M. S., Tanaka, K., Iwamoto, D. V., Calderwood, D. A., Svitkina, T. M., & Schwartz, M. A. (2019). Filamin A mediates isotropic distribution of applied force across the actin network. Journal of Cell Biology, 218(8), 2481–2491. https://doi.org/10.1083/jcb.201901086

55. Lehmann, R., & Tautz, D. (1994). In Situ Hybridization to RNA. Methods in Cell Biology, 44(C), 575–596. https://doi.org/10.1016/S0091-679X(08)60933-4

56. Lemaitre, B., & Hoffmann, J. (2007). The Host Defense of *Drosophila melanogaster*. Annual Review of Immunology, 25(1), 697–743. https://doi.org/10.1146/annurev.immunol.25.022106.141615

57. Lesch, C., Jo, J., Wu, Y., Fish, G. S., & Galko, M. J. (2010). A Targeted *UAS-RNAi* Screen in Drosophila Larvae Identifies Wound Closure Genes Regulating Distinct Cellular Processes. Genetics, 186(3), 943–957. https://doi.org/10.1534/genetics.110.121822

58. Lian, G., Dettenhofer, M., Lu, J., Downing, M., Chenn, A., Wong, T., & Sheen, V. (2016). Filamin A- and formin 2-dependent endocytosis regulates proliferation via the canonical wnt pathway. Development (Cambridge), 143(23). https://doi.org/10.1242/dev.139295

59. Linder, M., Glitzner, E., Srivatsa, S., Bakiri, L., Matsuoka, K., Shahrouzi, P., … Sibilia, M. (2018). EGFR is required for FOS dependent bone tumor development via RSK2/CREB signaling. EMBO Molecular Medicine, 10(11). https://doi.org/10.15252/emmm.201809408

60. Luster, A. D., Alon, R., & von Andrian, U. H. (2005). Immune cell migration in inflammation: Present and future therapeutic targets. Nature Immunology, 6(12), 1182–1190. https://doi.org/10.1038/ni1275

61. Makhijani, K., Alexander, B., Tanaka, T., Rulfson, E., & Brückner, K. (2011). The peripheral nervous system supports blood cell homing and survival in the Drosophila larva. Development, 138(24), 5379–91. https://doi.org/10.1242/dev.067322

62. Matsubayashi, Y., Louani, A., Dragu, A., Sánchez-Sánchez, B. J., Serna-Morales, E., Yolland, L., … Stramer, A. M. (2017). A Moving Source of Matrix Components Is Essential for De Novo Basement Membrane Formation. Current Biology, 27(22), 3526–3534.e4. https://doi.org/10.1016/j.cub.2017.10.001

63. Min, P. I., Spurney, R. F., Qisheng, T. U., Hinson, T., & Darryl Quarles, L. (2002). Calcium-sensing receptor activation of Rho involves filamin and rho-guanine nucleotide exchange factor. Endocrinology, 143(10), 3830–3838. https://doi.org/10.1210/en.2002-220240

64. Mitchison, T.J., and Cramer, L.P. (1996) Actin-based cell motility and cell locomotion. Cell 84(3)371–9). https://doi.org/10.1016/s0092-8864(00)81281-7.

65. Muñoz-Alarcón, A., Pavlovic, M., Wismar, J., Schmitt, B., Eriksson, M., Kylsten, P., & Dushay, M. S. (2007). Characterization of lamin mutation phenotypes in Drosophila and comparison to human laminopathies. PloS One, 2(6). https://doi.org/10.1371/journal.pone.0000532

66. Ohta, Y., Suzuki, N., Nakamura, S., Hartwig, J. H., & Stossel, T. P. (1999). The small GTPase RaIA targets filamin to induce filopodia. Proceedings of the National Academy of Sciences of the United States of America, 96(5), 2122–2128. https://doi.org/10.1073/pnas.96.5.2122

67. Paluch, E. K., Aspalter, I. M., & Sixt, M. (2016). Focal Adhesion–Independent Cell Migration. Annual Review of Cell and Developmental Biology, 32(1), 469–490. https://doi.org/10.1146/annurev-cellbio-111315-125341

68. Perez-Hernandez, D., Gutiérrez-Vázquez, C., Jorge, I., López-Martín, S., Ursa, A., Sánchez-Madrid, F., … Yañez-Mó, M. (2013). The intracellular interactome of tetraspanin-enriched microdomains reveals their function as sorting machineries toward exosomes. Journal of Biological Chemistry, 288(17), 11649–11661. https://doi.org/10.1074/jbc.M112.445304

69. Perkins, L. A., Holderbaum, L., Tao, R., Hu, Y., Sopko, R., McCall, K., … Perrimon, N. (2015). The transgenic RNAi project at Harvard medical school: Resources and validation. Genetics, 201(3), 843–852. https://doi.org/10.1534/genetics.115.180208

70. Popowicz, G. M., Schleicher, M., Noegel, A. A., & Holak, T. A. (2006). Filamins: promiscuous organizers of the cytoskeleton. Trends in Biochemical Sciences, 31(7), 411–419. https://doi.org/10.1016/j.tibs.2006.05.006

71. Raab, M., Gentili, M., de Belly, H., Thiam, H.R., Vargas, P., Jimenez, A.J., Lautenschlaeger, F., Voituriez, R., Lennon-Duménil, A.M., Manel, N, Piel, M. (2016). ESCRT III repairs nuclear envelope ruptures during cell migration to limit DNA damage and cell death. Science, 352(6283), 359–62. https://doi.org/10.1126/science.aad7611

72. Ratheesh, A., Belyaeva, V., & Siekhaus, D. E. (2015). Drosophila immune cell migration and adhesion during embryonic development and larval immune responses. Current Opinion in Cell Biology, 36, 71–79. https://doi.org/10.1016/j.ceb.2015.07.003

73. Ratheesh, A., Biebl, J., Vesela, J., Smutny, M., Papusheva, E., Krens, S. F. G., … Siekhaus, D. E. (2018). Drosophila TNF Modulates Tissue Tension in the Embryo to Facilitate Macrophage Invasive Migration. Developmental Cell, 45(3), 331–346.e7. https://doi.org/10.1016/j.devcel.2018.04.002

74. Razinia, Z., Mäkelä, T., Ylänne, J., & Calderwood, D. A. (2012). Filamins in Mechanosensing and Signaling. Annual Review of Biophysics, 41(1), 227–246. https://doi.org/10.1146/annurev-biophys-050511-102252

75. Riesgo-Escovar, J. R., & Hafen, E. (1997). Common and distinct roles of DFos and DJun during Drosophila development. Science (New York, N.Y.), 278(5338), 669–672. https://doi.org/10.1126/science.278.5338.669

76. Riveline, D., Zamir, E., Balaban, N. Q., Schwarz, U. S., Ishizaki, T., Narumiya, S., … Bershadsky, A. D. (2001). Focal contacts as mechanosensors: Externally applied local mechanical force induces growth of focal contacts by an mDia1-dependent and ROCK-independent mechanism. Journal of Cell Biology, 153(6), 1175–1185. https://doi.org/10.1083/jcb.153.6.1175

77. Rose, R., Weyand, M., Lammers, M., Ishizaki, T., Ahmadian, M. R., & Wittinghofer, A. (2005). Structural and mechanistic insights into the interaction between Rho and mammalian Dia. Nature, 435(7041), 513–518. https://doi.org/10.1038/nature03604

78. Rousso, T., Shewan, A. M., Mostov, K. E., Schejter, E. D., & Shilo, B. Z. (2013). Apical targeting of the formin diaphanous in Drosophila tubular epithelia. ELife, 2013(2). https://doi.org/10.7554/eLife.00666

79. Sánchez-Sánchez, B. J., Urbano, J. M., Comber, K., Dragu, A., Wood, W., Stramer, B., & Martín-Bermudo, M. D. (2017). Drosophila Embryonic Hemocytes Produce Laminins to Strengthen Migratory Response. Cell Reports, 21(6), 1461–1470. https://doi.org/10.1016/j.celrep.2017.10.047

80. Seth, A., Otomo, C., & Rosen, M. K. (2006). Autoinhibition regulates cellular localization and actin assembly activity of the diaphanous-related formins FRLα and mDia1. Journal of Cell Biology, 174(5), 701–713. https://doi.org/10.1083/jcb.200605006

81. Sharma, P. & Allison, J.P. (2015) The future of immune checkpoint therapy. Science 348(6230), 56–61. https://doi.org/10.1126/science.aaa8172.

82. Shigeta, M., Sanzen, N., Ozawa, M., Gu, J., Hasegawa, H., & Sekiguchi, K. (2003). CD151 regulates epithelial cell-cell adhesion through PKC- and Cdc42-dependent actin cytoskeletal reorganization. Journal of Cell Biology, 163(1), 165–176. https://doi.org/10.1083/jcb.200301075

83. Siekhaus, D., Haesemeyer, M., Moffitt, O., & Lehmann, R. (2010). RhoL controls invasion and Rap1 localization during immune cell transmigration in Drosophila. Nature Cell Biology, 12(6), 605–610. https://doi.org/10.1038/ncb2063

84. Sit, S. T., & Manser, E. (2011). Rho GTPases and their role in organizing the actin cytoskeleton. Journal of Cell Science, 124(5), 679–683. https://doi.org/10.1242/jcs.064964

85. Smutny, M., Ákos, Z., Grigolon, S., Shamipour, S., Ruprecht, V., Čapek, D., … Heisenberg, C. P. (2017). Friction forces position the neural anlage. Nature Cell Biology, 19(4), 306–317. https://doi.org/10.1038/ncb3492

86. Sokol, N. S., & Cooley, L. (2003). Drosophila filamin is required for follicle cell motility during oogenesis. Developmental Biology, 260(1), 260–272. https://doi.org/10.1016/S0012-1606(03)00248-3

87. Somogyi, K., & Rørth, P. (2004). Evidence for tension-based regulation of Drosophila MAL and SRF during invasive cell migration. Developmental Cell, 7(1), 85–93. https://doi.org/10.1016/j.devcel.2004.05.020

88. Stossel, T. P., Condeelis, J., Cooley, L., Hartwig, J. H., Noegel, A., Schleicher, M., & Shapiro, S. S. (2001). Filamins as integrators of cell mechanics and signalling. Nature Reviews Molecular Cell Biology, 2(2), 138–145. https://doi.org/10.1038/35052082

89. Szabo, P.A., Miron, M., & Farber, D.L. (2019). Location, location, location: Tissue residence memory T cells in mice and humans. Science Immunology 4(34), https://doi.org/10.1126/scieimunol.aas9673

90. Szalóki, N., Krieger, J. W., Komáromi, I., Tóth, K., & Vámosi, G. (2015). Evidence for Homodimerization of the c-Fos Transcription Factor in Live Cells Revealed by Fluorescence Microscopy and Computer Modeling. Molecular and Cellular Biology, 35(21). https://doi.org/10.1128/mcb.00346-15

91. Tejera, E., Rocha-Perugini, V., López-Martín, S., rez-Hernández, D., Bachir, A. I., Horwitz, A. R., … Yáñez-Mo, M. (2013). CD81 regulates cell migration through its association with Rac GTPase. Molecular Biology of the Cell, 24(3), 261–273. https://doi.org/10.1091/mbc.E12-09-0642

92. Termini, C. M., & Gillette, J. M. (2017). Tetraspanins Function as Regulators of Cellular Signaling. Frontiers in Cell and Developmental Biology, 5, 34. https://doi.org/10.3389/fcell.2017.00034

93. Theret, M., Mounier, R., & Rossi, F. (2019). The origins and non-canonical functions of macrophages in development and regeneration. Development, 146(9). https://doi.org/10.1242/dev.156000

94. Thiam, H., Vargas, P., Carpi, N., Crespo, C.L., Raab, M., Terriac, E., King, M.C., Jacobelli, J., Alberts, A.S., Stradal, T., Lennon-Dumenil, A., Piel, M. (2016). Perinuclear Arp2/3-driven actin polymerization enables nuclear deformation to facilitate cell migration through complex environments. Stremmel:10997, https://doi.org/10.1038/ncomms10997

95. Thurmond, J., Goodman, J. L., Strelets, V. B., Attrill, H., Gramates, L. S., Marygold, S. J., … Baker, P. (2019). FlyBase 2.0: The next generation. Nucleic Acids Research, 47(D1). https://doi.org/10.1093/nar/gky1003

96. Tomancak, P., Beaton, A., Weiszmann, R., Kwan, E., Shu, S. Q., Lewis, S. E., … Rubin, G. M. (2002). Systematic determination of patterns of gene expression during Drosophila embryogenesis. Genome Biology, 3(12). https://doi.org/10.1186/gb-2002-3-12-research0088

97. Tomancak, P., Berman, B. P., Beaton, A., Weiszmann, R., Kwan, E., Hartenstein, V., … Rubin, G. M. (2007). Global analysis of patterns of gene expression during Drosophila embryogenesis. Genome Biology, 8(7). https://doi.org/10.1186/gb-2007-8-7-r145

98. Tseng, Y., An, K. M., Esue, O., & Wirtz, D. (2004). The Bimodal Role of Filamin in Controlling the Architecture and Mechanics of F-actin Networks. Journal of Biological Chemistry, 279(3), 1819–1826. https://doi.org/10.1074/jbc.M306090200

99. Uhlirova, M., & Bohmann, D. (2006). JNK- and Fos-regulated Mmp1 expression cooperates with Ras to induce invasive tumors in Drosophila. EMBO Journal, 25(22), 5294–5304. https://doi.org/10.1038/sj.emboj.7601401

100. Vadlamudi, R. K., Li, F., Adam, L., Nguyen, D., Ohta, Y., Stossel, T. P., & Kumar, R. (2002). Filamin is essential in actin cytoskeletal assembly mediated by p21-activated kinase 1. Nature Cell Biology, 4(9), 681–690. https://doi.org/10.1038/ncb838

101. Valoskova, K., Biebl, J., Roblek, M., Emtenani, S., Gyoergy, A., Misova, M., … Siekhaus, D. E. (2019). A conserved major facilitator superfamily member orchestrates a subset of O-glycosylation to aid macrophage tissue invasion. ELife, 8. https://doi.org/10.7554/eLife.41801

102. Vetter, I.R., & Wittinghofer, A. (2001). The guanine nucleotide-binding switch in three dimensions. Science 294(5545),1299–1304. https://doi.org/10.1126/science.1062023

103. Warner, S. J., & Longmore, G. D. (2009). Cdc42 antagonizes Rho1 activity at adherens junctions to limit epithelial cell apical tension. Journal of Cell Biology, 187(1), 119–133. https://doi.org/10.1083/jcb.200906047

104. Weavers, H., Evans, I. R., Martin, P., & Wood, W. (2016). Corpse Engulfment Generates a Molecular Memory that Primes the Macrophage Inflammatory Response. Cell. https://doi.org/10.1016/j.cell.2016.04.049

105. Williams, M. J., Habayeb, M. S., & Hultmark, D. (2007). Reciprocal regulation of Rac1 and Rho1 in Drosophila circulating immune surveillance cells. Journal of Cell Science, 120(3), 502–511. https://doi.org/10.1242/jcs.03341

106. Wintner, O., Hirsch-Attas, N., Schlossberg, M., Brofman, F., Friedman, R., Kupervaser, M., … Buxboim, A. (2020). A Unified Linear Viscoelastic Model of the Cell Nucleus Defines the Mechanical Contributions of Lamins and Chromatin. Advanced Science, 7(8). https://doi.org/10.1002/advs.201901222

107. Wood, W., Faria, C., & Jacinto, A. (2006). Distinct mechanisms regulate hemocyte chemotaxis during development and wound healing in Drosophila melanogaster. Journal of Cell Biology, 173(3). https://doi.org/10.1083/jcb.200508161

108. Yáñez-Mó, M., Barreiro, O., Gordon-Alonso, M., Sala-Valdés, M., & Sánchez-Madrid, F. (2009). Tetraspanin-enriched microdomains: a functional unit in cell plasma membranes. Trends in Cell Biology, 19(9), 434–446. https://doi.org/10.1016/j.tcb.2009.06.004

109. Yeung, L., Hickey, M.J., & Wright, M.D. (2018) The many and varied roles of tetraspanins in immune cell recrutment and migration. Frontiers in Immunology.9:1644. https:// doi: 10.3389/fimmu.2018.01644.

110. Zeitlinger, J., Kockel, L., Peverali, F. A., Jackson, D. B., Mlodzik, M., & Bohmann, D. (1997). Defective dorsal closure and loss of epidermal decapentaplegic expression in Drosophila fos mutants. EMBO Journal, 16(24), 7393–7401. https://doi.org/10.1093/emboj/16.24.7393

111. Zhang, X. A., Bontrager, A. L., & Hemler, M. E. (2001). Transmembrane-4 Superfamily Proteins Associate with Activated Protein Kinase C (PKC) and Link PKC to Specific β1 Integrins. Journal of Biological Chemistry, 276(27), 25005–25013. https://doi.org/10.1074/jbc.M102156200

112. Zhou, J., Kim, H. Y., & Davidson, L. A. (2009). Actomyosin stiffens the vertebrate embryo during crucial stages of elongation and neural tube closure. Development, 136(4), 677–688. https://doi.org/10.1242/dev.026211

113. Zimmerman, B., Kelly, B., McMillan, B. J., Seegar, T. C. M., Dror, R. O., Kruse, A. C., & Blacklow, S. C. (2016). Crystal Structure of a Full-Length Human Tetraspanin Reveals a Cholesterol-Binding Pocket. Cell, 167(4). https://doi.org/10.1016/j.cell.2016.09.056

114. Zimmerman, B., Kelly, B., McMillan, B., Seegar, T., Kruse, A., & Blacklow, S. (2016). Crystal Structure of Human Tetraspanin CD81 Reveals a Conserved Intramembrane Binding Cavity. Cell, 167(4), 1041–1051. https://doi.org/10.1016/j.cell.2016.09.056

115. Zwerger, M., Jaalouk, D. E., Lombardi, M. L., Isermann, P., Mauermann, M., Dialynas, G., … Lammerding, J. (2013). Myopathic lamin mutations impair nuclear stability in cells and tissue and disrupt nucleo-cytoskeletal coupling. Human Molecular Genetics, 22(12), 2335–2349. https://doi.org/10.109

