## Supplementary figures and images for "Cortical actin properties controlled by *Drosophila* Fos aid macrophage infiltration against surrounding tissue resistance"

### S1 Figure

#### Belayeva et al, Figure 1- supplement figure 1

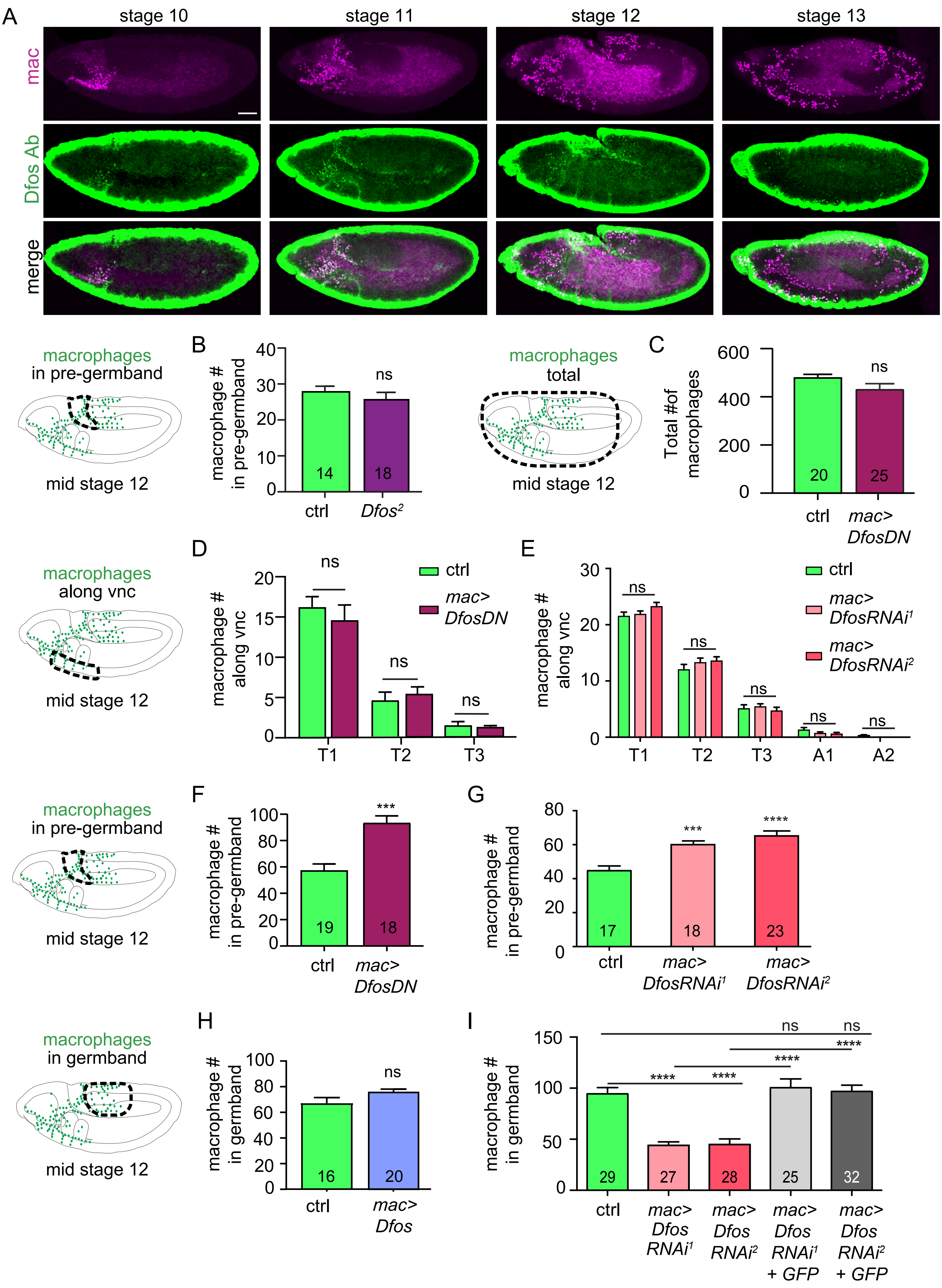

### S2 Figure

#### Belyaeva et al, Figure 2- supplement figure 1

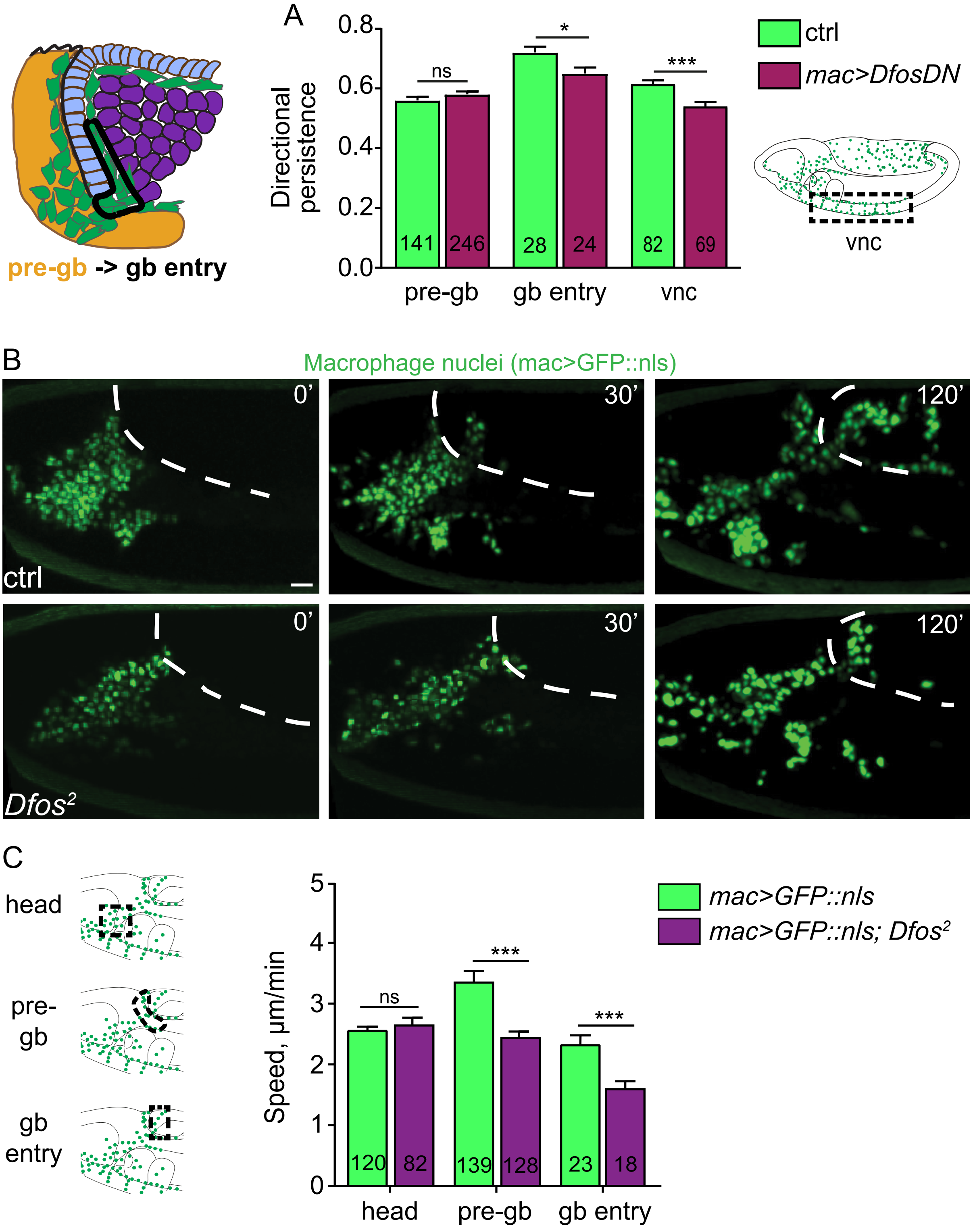

### S3 Figure

#### Belyaeva et al, Figure 3- supplement figure 1

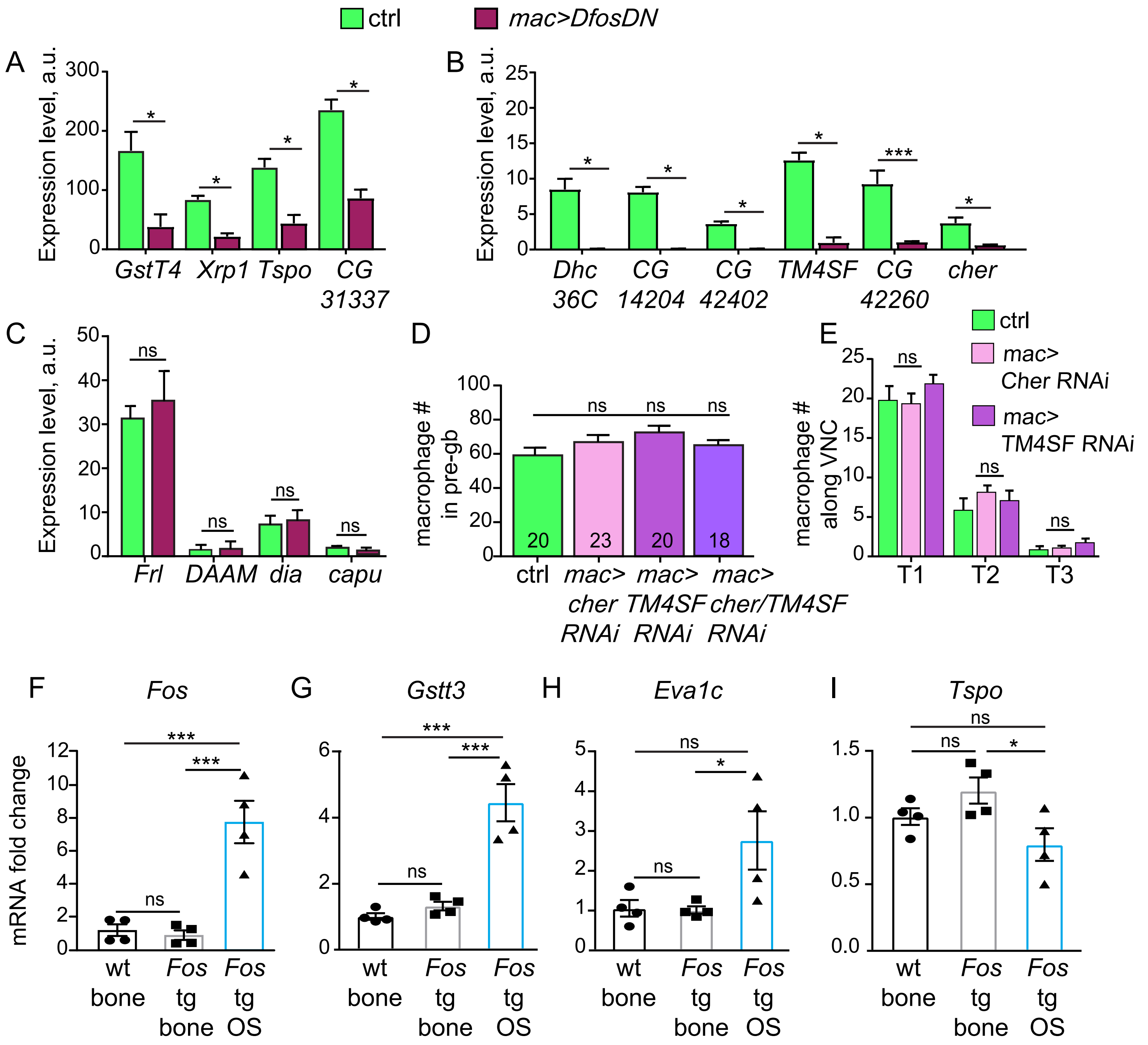

### S4 Figure

#### Belyaeva et al, Figure 4- supplement figure 1

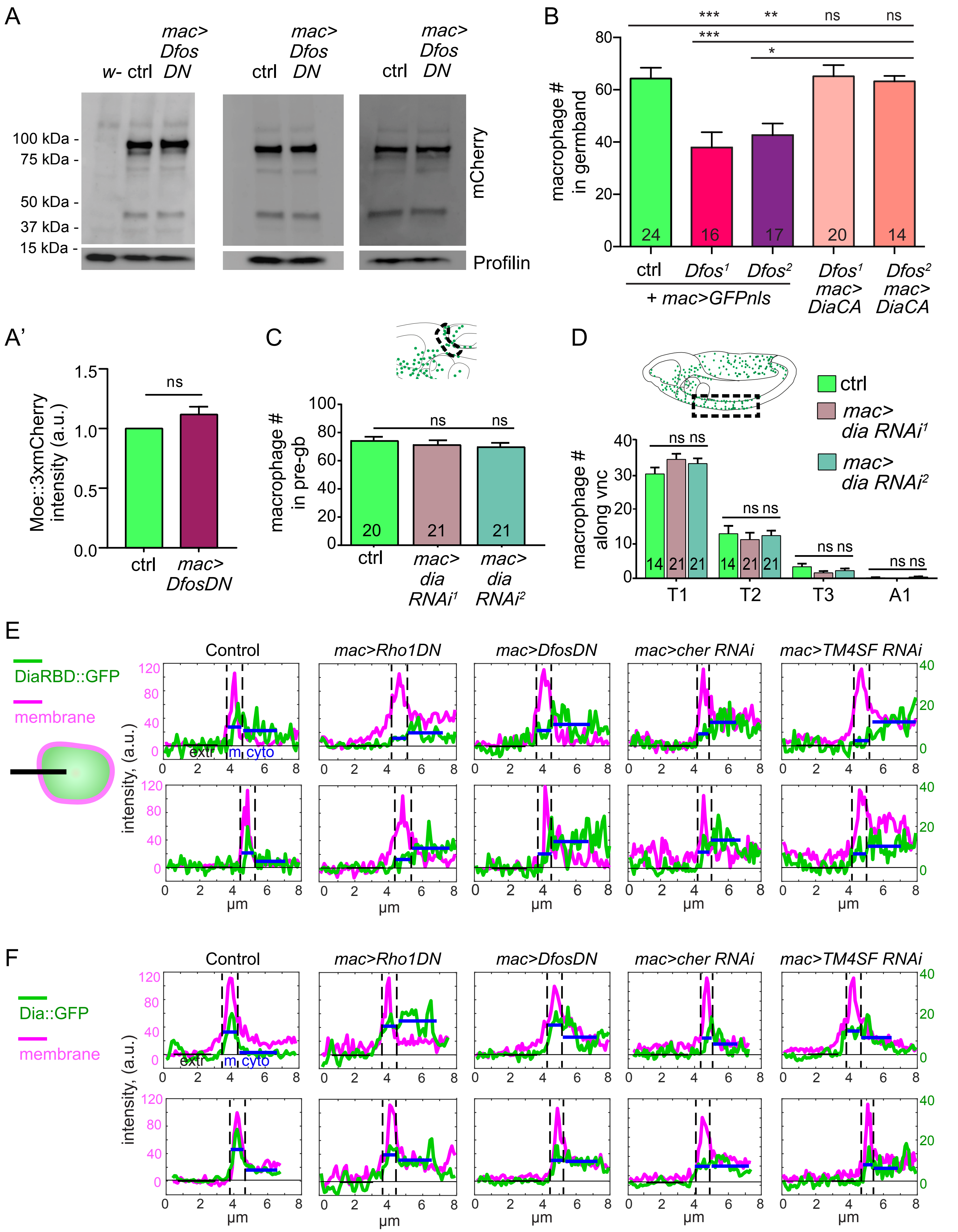

### S5 Figure

Belyaeva et al, Figure 5- supplement figure 1

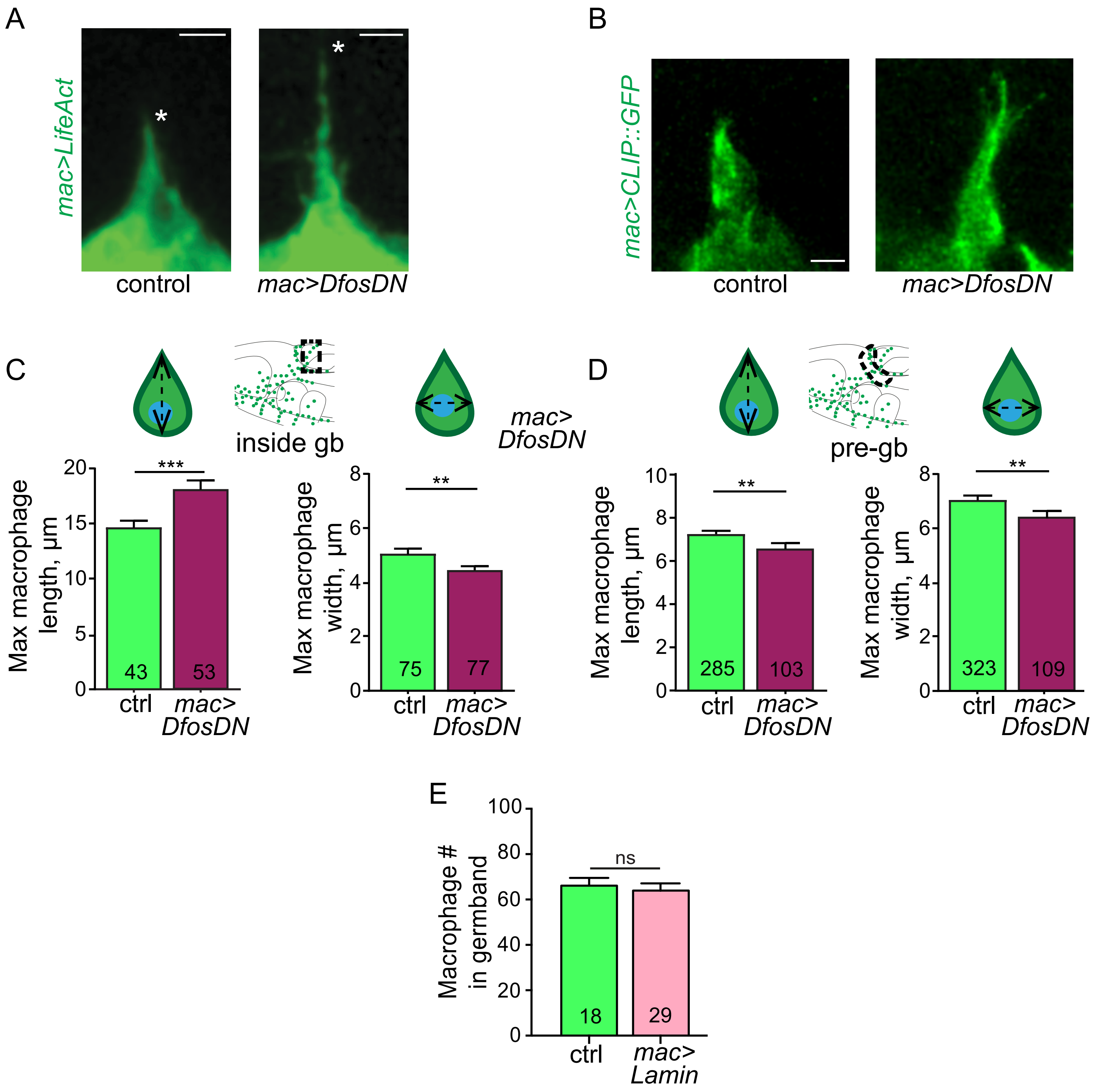
