## Supplementary material for "Cortical actin properties controlled by *Drosophila* Fos aid macrophage infiltration against surrounding tissue resistance": S6 Figure

### Belyaeva et al, Figure 6- supplement figure 1

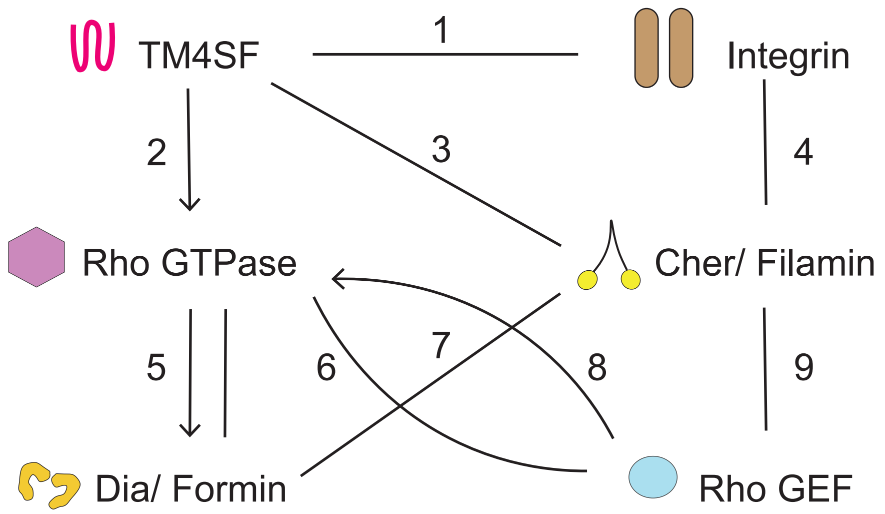


**Figure 6—figure supplement 1. Model of protein interactions at the macrophage cortex**

Proposed protein interactions at the cell cortex in wildtype macrophages during germband infiltration as shown in Figure 6. Direct binding between two proteins is indicated by a line, signalling between the interaction partners is represented as an arrow. These interactions and the resulting model in Figure 6 are based on the papers listed at the end of this legend next to the corresponding number shown for each particular linkage. The Tetraspanin TM4SF can cluster adhesion receptors such as Integrins at the membrane and lead to the recruitment and activation of Rho GTPases. Rho GTPases can bind and activate the formin Dia leading to F-actin polymerization. In addition, Integrin can bind filamins (Cher), which can bind to and thereby recruit RhoGEF to the membrane. Rho GEFs can in turn bind to and activate Rho GTPases. References for listed interactions: **1, Tetraspanins-Integrin**) Berditchevski et al., 2001; Berditchevski & Odintsova, 1999; Fitter et al., 1999; Hemler et al., 1996; Kazarov et al, 2002; Levy & Shoham, 2005; Sterk et al., 2002; Tachibana et al., 1997; Termini & Gillette, 2017; Winterwood et al., 2006; Yáñez-Mó et al., 2009; X. H. Yang et al., 2008; X. Yang et al., 2004; Yauch et al., 1998; Yaucht et al., 2000; X. A. Zhang et al., 2001. **2, Tetraspanins-Rho GTPases**) Delaguillaumie et al., 2002; Herr et al.,, 2014; Hong et al., 2012; Imhof et al., 2008; Jones et al., 2016; W. M. Liu et al., 2012; Novitskaya et al., 2014; Shigeta et al., 2003; Sit & Manser, 2011; Tejera et al., 2013; F. Zhang et al., 2011. **3, Tetraspanins-Filamins**) Brzozowski et al., 2018; Perez-Hernandez et al., 2013. **4, Integrin-Filamins**) Chen et al., 2009; Das et al., 2011; Ehrlicher et al., 2011; Kumar et al., 2019; Razinia et al., 2012. **5, Rho GTPases-Formins**) Alberts, 2001; Bechtold et al., 2014; Eisenmann et al., 2007; Großhans et al., 2005; Homem & Peifer, 2009; Kühn & Geyer, 2014; Li & Higgs, 2003, 2005; Massarwa et al., 2009; Rose et al., 2005; Rousso et al., 2013; Seth et al., 2006; Watanabe et al., 1999; Watanabe et al., 1997; Williams et al., 2007. **6 & 8, Rho GEF-Rho GTPases**) García-Mata & Burridge, 2007; Goicoechea et al., 2014; Lawson & Burridge, 2014; Rossman et al., 2005) **7, Formins-Filamins**) (Hu et al., 2014; Lian et al., 2016; Luo et al., 2013) **9, Filamins-RhoGEFs**) (Bellanger et al., 2000; Bourguignon et al., 2006; Min et al., 2002; Van Rijssel et al., 2012)
