## Supplementary material for "Cortical actin properties controlled by *Drosophila* Fos aid macrophage infiltration against surrounding tissue resistance": Table S1

**S1 Table. Comparison of transcription factor (TF) mRNA expression in macrophages at stages 11-12 and 13-16**

Macrophages acquire different TF profiles at different stages of embryonic development (based on Hammonds et al., 2013). TFs expressed in the macrophages only at stages 11-12 are highlighted in green. TFs expressed in macrophages only at stages 13-16 are highlighted in blue. Function is annotated only for TFs expressed in macrophages at stages 11-12.

| Stages 11-12 | Stages 13-16 | Function |
|---|---|---|
| *brk* | | Dpp-related, development |
| *Bteb2* | *Bteb2* | |
| *c15* | *c15* | |
| *croc* | | Early embryo patterning |
| | *Dif* | |
| | *dm* | |
| *foxo* | *foxo* | |
| *gcm* | *gcm* | |
| | *gcm2* | |
| *ham* | | Neuronal cell fate |
| *Dfos* | | Cell polarity, wound healing, development, cell cycle |
| *kn* | | Early embryo patterning |
| *lz* | *lz* | |
| *Mad* | *Mad* | |
| *MafS* | | Transcription |
| *MTF1* | *MTF-1* | |
| *odd* | *odd* | |
| | *Pdp1* | |
| | *pros* | |
| | *Rel* | |
| *six4* | | Mesodermal patterning, gonad development |
| *slp1* | *slp1* | |
| *srp* | *srp* | |
| *svp* | *svp* | |
| *tj* | | Development |
| *topi* | | Spermatid differentiation |
| | *Usf* | |
| | *ush* | |

| | | |
|---|---|---|
| *vri* | | Cell growth, tracheal development, circadian rhythm |
| *zfh1* | *zfh1* | |
| *CG17801* | | Unknown |
| *CG17802* | | Unknown |
| *CG33213* | *CG33213* | |
| | *CG30431* | |
| | *CG11071* | |
| | *CG8145* | |
| | *CG9932* | |
