## Supplementary material for "Cortical actin properties controlled by *Drosophila* Fos aid macrophage infiltration against surrounding tissue resistance": Table S2

**S2 Table. Genes upregulated in macrophages expressing DfosDN**

Genes are ordered according to the adjusted p-value from the RNA-Sequencing.

The murine ortholog with the top score in UniProt BLAST is shown in the table.

| Gene | Adjusted p-value | Possible function | Mouse ortholog, Identity |
|---|---|---|---|
| *CG1673* | 0.0002 | Transamination of branched-chain-amino-acids | Branched-chain-amino-acid aminotransferase, 53.9% |
| *Hsp70Ab* | 0.0007 | Chaperone, protein folding | Heat shock 70 kDa protein 1B, 72.2% |
| *l(1)G0469* | 0.01 | Unknown | SKI/DACH domain-containing protein 1, 44.8% |
| *Hsp68* | 0.02 | Unfolded protein binding | Heat shock 70 kDa protein 1B, 72.2% |
| *CG13321* | 0.03 | Unknown | Tensin-2, 34.1% |
| *Hsp70Aa* | 0.03 | Chaperone, protein folding | Heat shock 70 kDa protein 1A, 76.3% |
| *Sug* | 0.03 | Transcription | Zinc finger protein GLIS2, 57.4% |
| *Hsp70Bc* | 0.03 | Chaperone, protein folding | Heat shock 70 kDa protein 1A, 76.6% |
| *CG6574* | 0.046 | Transport of folate | Thiamine transporter 1, 37.3% |
